# Transcriptomic and proteomic dynamics of ovarian follicle group culture resemble *in vivo* folliculogenesis

**DOI:** 10.1101/2025.09.12.675858

**Authors:** Andrea S. K. Jones, D. Ford Hannum, Taylor Schissel, Jordan H. Machlin, Vasantha Padmanabhan, Jun Z. Li, Ariella Shikanov

**Affiliations:** Department of Biomedical Engineering, University of Michigan, Ann Arbor, MI; Department of Computational Medicine and Bioinformatics, University of Michigan, Ann Arbor, MI; Cellular and Molecular Biology Program, University of Michigan, Ann Arbor, MI; Department of Obstetrics and Gynecology, University of Michigan, Ann Arbor, MI; Department of Pediatrics and Communicable Diseases, University of Michigan, Ann Arbor, MI; Department of Molecular and Integrative Physiology, University of Michigan, Ann Arbor, MI; Department of Molecular Genetics and Genome Sciences, University of Oklahoma, Oklahoma City, OK

## Abstract

The prohibitively low yield of fertilizable oocytes obtained from cultured ovarian follicles limits clinical translation of *in vitro* follicle maturation for fertility preservation. This is in part due to an incomplete understanding of the process of follicle development. Previous work has demonstrated that group culture of primary murine follicles had a synergistic effect on growth and maturation in contrast to single follicles, but mechanisms remained unknown. Here, we cultured primary follicles in groups of 5 (5X) or 10 (10x) for twelve days, separated the somatic cells from oocytes, and analyzed the temporal transcriptional signatures every two days. In total, 13,461 genes in somatic cells and 10,091 genes in oocytes were computationally sorted into ten temporally distinct gene expression patterns. The somatic cell temporal gene expression patterns showed strong concordance with the granulosa and theca cell markers reported in a recent single-cell whole ovary RNA sequencing study. Importantly, canonical markers of steroidogenesis in cultured follicles followed expected trajectories of decreasing *Amh* expression and increasing *Inhba, Inhbb, Cyp11a1, Cyp17a1, Cyp19a1, Lhcgr,* and *Fshr* over the culture period. Furthermore, when comparing the 10X and 5X culture groups, we identified 306 and 14 differentially expressed genes in somatic cells and oocytes, respectively. Shotgun proteomics data was aligned with the somatic cell transcriptomic data and identified four L-R pairs that were differentially expressed between the two conditions. These comprehensive datasets uncovered temporal dynamics of *in vitro* folliculogenesis in a compartment-specific manner, serving as a valuable resource for optimizing future follicle culture systems for fertility preservation.

## INTRODUCTION

The ovarian follicle is the multicellular functional unit of the ovary that is the source of fertility. Each follicle contains an oocyte and two types of hormone-producing somatic cells, granulosa and theca cells, which differentiate over the course of folliculogenesis into a number of subtypes with distinct functions. Follicle development, or folliculogenesis, is a coordinated effort by the growing oocyte and proliferating granulosa and theca cells, which perform critical roles in steroid hormone production and oocyte maturation. At the earliest stage of maturation, the primordial follicle is maintained in a state of arrest and is comprised of the oocyte surrounded by a single layer of squamous granulosa cells. Follicles are non-renewable and their number declines over time as the majority of follicles undergo a natural degenerative process called atresia, while a select number of follicles progress to ovulation during the reproductive lifespan. This loss of the follicular reserve is accelerated in patients who undergo anti-cancer treatments including chemotherapy and radiation that cause irreversible follicle loss and can result in primary ovarian insufficiency and infertility. Therefore, patients often seek fertility preservation prior to treatment via oocyte retrieval followed by oocyte or embryo cryopreservation. However, this approach is problematic for patients who have aggressive cancers and cannot delay treatment, or for prepubertal patients with immature follciles^1–5^. Alternatively, isolation of immature early-stage ovarian follicles, which constitute the majority of the ovarian reserve, followed by follicle maturation *in vitro,* poses as a promising approach to fertility preservation that does not interfere with anti-cancer treatment.

Efforts to culture early-stage (primordial and primary) follicles have had limited success due to poor replication of the ovarian microenvironment *in vitro*. *In vivo,* follicles rely on a complex paracrine milieu for proper maturation^6–8^, and the surrounding extracellular matrix provides structural support to their 3D architecture. Biomimetic hydrogel systems are commonly employed to provide mechanical support to growing follicles *in vitro*. Likewise, follicles have been cultured in groups to enrich for the follicle-derived, currently uncharacterized soluble cues that support maturation. Recently, we and others reported that group culture of primary follicles significantly improved survival and growth when compared to individually cultured follicles, with groups of ten (10X) being more successful than groups of five (5X)^9–11^. To identify the mechanisms driving this improved folliculogenesis, we previously used microarray-based gene expression analysis of somatic cells isolated from 5X and 10X follicles throughout culture. This analysis revealed differences in the transcriptomic profile with dynamic expression of genes related to prolactin signaling (10X: increasing across culture, 5X: decreasing across culture) and angiogenesis (10X: decreasing across culture, 5X: increasing followed by decreasing)^11^. However, separate transcriptomic analysis of oocytes isolated from cultured follicles was not performed due to technical limitations, leaving a gap in our understanding of how bidirectional communication between the oocyte and somatic cells impacts folliculogenesis.

Here, we expanded on our microarray-based analysis of the transcriptomes of 5X and 10X follicles using a more robust RNA sequencing (RNA-seq) approach and separately analyzed the follicular somatic cells and corresponding oocytes as paired samples. We collected transcriptome data over six time points spanning the 12-day culture period (every other day from Day 0 to Day 12) to identify dynamic changes during culture, and we compared temporal dynamics between the 5X and 10X conditions. We uncovered a series of ten temporally-related gene expression patterns for the 13,461 genes that were differentially expressed over time (DET) in somatic cells and the 10,091 DET genes in oocytes. Within the somatic cells, these patterns matched the granulosa and theca cell markers reported in a recent study from follicles that developed naturally *in vivo.*^12^ Comparison revealed that there are more genes differentially expressed across 5X and 10X conditions (DEC) in somatic cells (306 genes) than oocytes (14 genes). We also performed ligand-receptor (L-R) analysis to identify oocyte-somatic cell interactions, finding more than a thousand temporally dynamic L-R gene pairs for each interaction type (Cell-Cell, Cell-Oocyte, Oocyte-Cell, and Oocyte-Oocyte) as well as 32 L-R DEC gene pairs. Finally, we used shotgun proteomics to validate and augment our transcriptome analysis, where we observed numerous points of agreement with the RNA-seq datasets and established additional proteins and biological processes associated with successful follicle development *in vitro*.

## RESULTS

### Growth and survival kinetics of longitudinal group follicle culture experiments

To profile the transcriptome dynamics of oocyte and somatic cell pairs of growing alginate-encapsulated primary murine follicles (5X and10X) over 12 days of culture, in 2-day intervals, and analyzed the transcriptome data (Fig. 1A,B). Survival rate across 5X and 10X conditions were similar, reaching 73% and 86% respectively on Day 12 (Fig. 1C). However, despite similar starting sizes on Day 0 (5X averaged by hydrogel: 102.6 µm, 10X averaged by hydrogel: 102.7 µm, p > 0.05; 5X by follicle: 102.1 µm, 10X by follicle: 102.3 µm, p > 0.05) (Fig. 1D), 10X follicles grew faster and larger (p=0.0013), reaching diameters of 301.6 ± 5.5 µm (mean ± SE) by Day 12, compared to 251.6 ± 10.6 µm for 5X follicles (Fig. 1E,F). Initial follicle diameter is reported in two ways: first, we show the average diameter of all follicles in a hydrogel, which demonstrates that each hydrogel contained follicles of similar starting sizes. Second, we show the diameter of each individual follicle and compare across conditions, showing that neither condition (5X and 10X) started off with an advantage of larger, later-stage follicles compared to the other.

**Figure 1:**
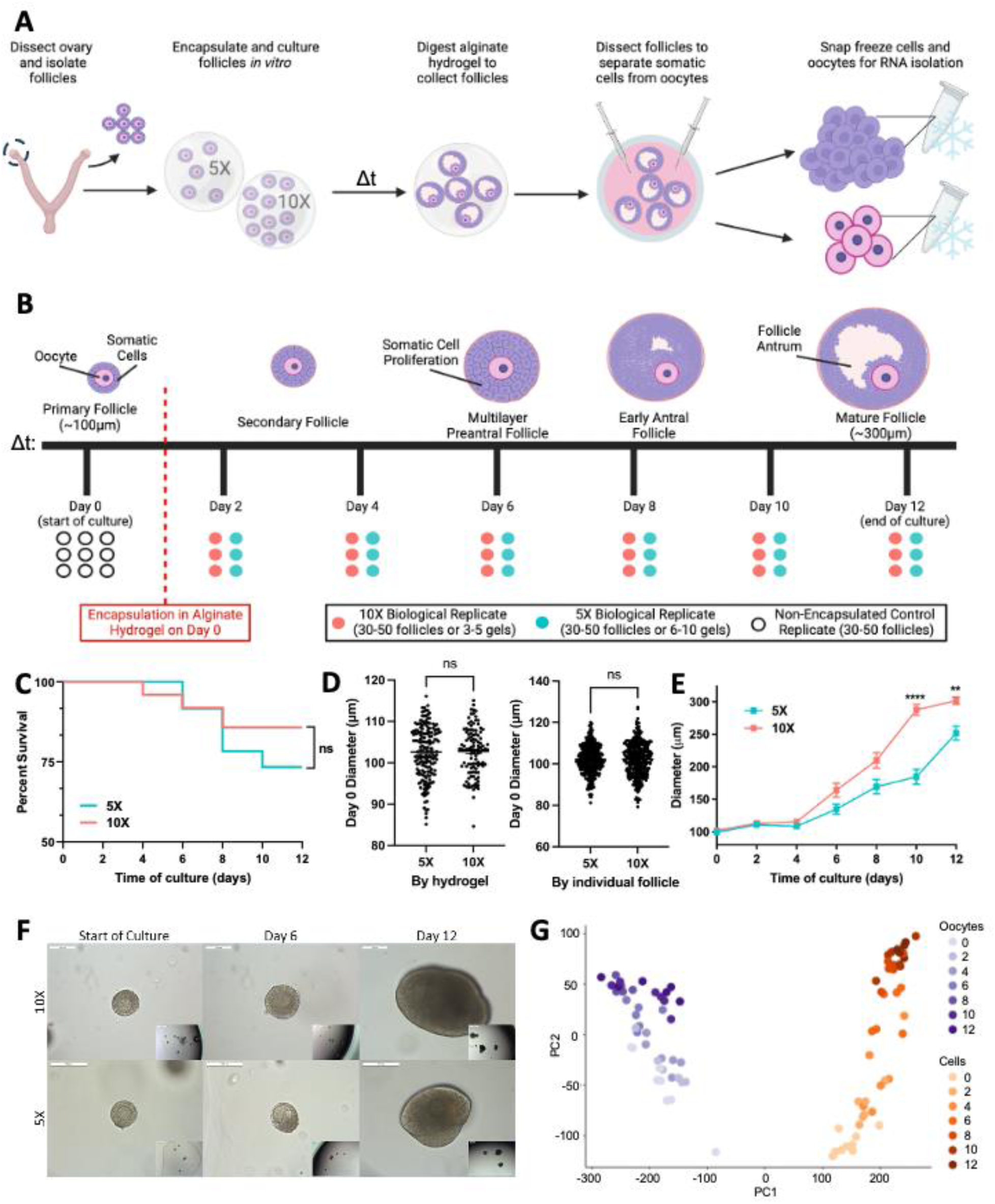
Ovarian follicles cultured in 10X outperform 5X culture. **A**) Schematic describing the experimental design, beginning with primary follicles cultured in alginate hydrogels in groups of 5 (5X) or 10 (10X). To collect samples for sequencing, hydrogels are enzymatically digested and the somatic cells are separated from their matched oocytes, **B)** Schematic describing the samples collected for sequencing. Over 12 days of culture, follicles were dissected at two-day intervals. The scheme was completed for somatic cells (n=45 samples) and oocytes (n=45 samples), **C**) Kaplan-Meier survival plots of follicles over duration of culture in 5X and 10X, Initial diameter of the ovarian follicles for RNA sequencing, by hydrogel (5X or 10X) (left), visualizing no statistical difference in starting diameters across conditions, **D**) Initial diameter of ovarian follicles for RNA sequencing by individual follicle (right), **E**) Growth rates of follicles over duration of culture in 5X and 10X, **F**) Bright field images of individual primary follicles encapsulated in alginate hydrogels in 5X/10X (scale bar = 100 µm, inset scale bar = 500 µm), and **G**) PC1-2 projection of oocyte and somatic cell RNA samples with color intensity correlating to the Day. *p < 0.05, **p < 0.01, ***p < 0.001, ****p < 0.0001. Figures 1A and 1B were generated using Biorender.com.

To establish a baseline transcriptional signature for Day 0, we enzymatically digested ovaries from mice 10-12 days old, selected primary stage follicles by diameter, separated the oocytes from the somatic cells via microdissection, and immediately froze them as Day 0 samples (n=9 oocyte and somatic cell pairs) (Fig. 1B, Supplemental Table 1). Separately, primary stage follicles were encapsulated and cultured in 5X or 10X groups in alginate for up to twelve days. Every two days, some of the alginate gels were digested with alginate lyase before separating the oocytes and somatic cells from the growing follicles for flash freezing, while the other cultures were maintained and "harvested" at subsequent time points. The study followed a 6-2-2-3 design, with samples representing six time points (Days 2, 4, 6, 8, 10, and 12), two cell types (oocytes, somatic cells), and two conditions (5X and 10X), with three technical replicates each, resulting in 72 samples for D2-12. The full set of 90 samples is represented in Fig. 1B and Supplemental Table 1. Each RNA sample was extracted from a pool of oocytes or somatic cells to obtain sufficient RNA, from 100 follicles on Day 0 and 30-70 follicles on subsequent days (Fig. 1B, Supplemental Table 1). After data processing and batch correction (**Methods**), RNA-seq data for the 90 samples showed marked separation between the oocyte and the somatic cell samples (Fig. 1G). Thus, we analyzed oocytes and somatic cells separately for the Day and Condition differences.

### Dynamics of somatic cell transcriptomes during culture

The somatic cell samples showed progressive transcriptional changes from D2 to D12, with limited differences between 5X and 10X, as shown by a general progression from left to right on a principal component plot (Fig. 2A), similar to our findings from microarray data^11^. Two-way ANOVA with Day, Condition (5X vs. 10X), and an Interaction term (**Methods**) revealed 13,461 genes with significant changes across the days of culture (differentially expressed over time, DET) (false discovery rate (FDR) < 0.05; same threshold also applies for Condition and Interaction effects), 306 genes with an overall difference between Conditions (differentially expressed across conditions, DEC), and 728 genes with a significant "Interaction", meaning they had differing temporal profiles between 5X and 10X. The two-way ANOVA results are in Supplemental File 1.

**Figure 2:**
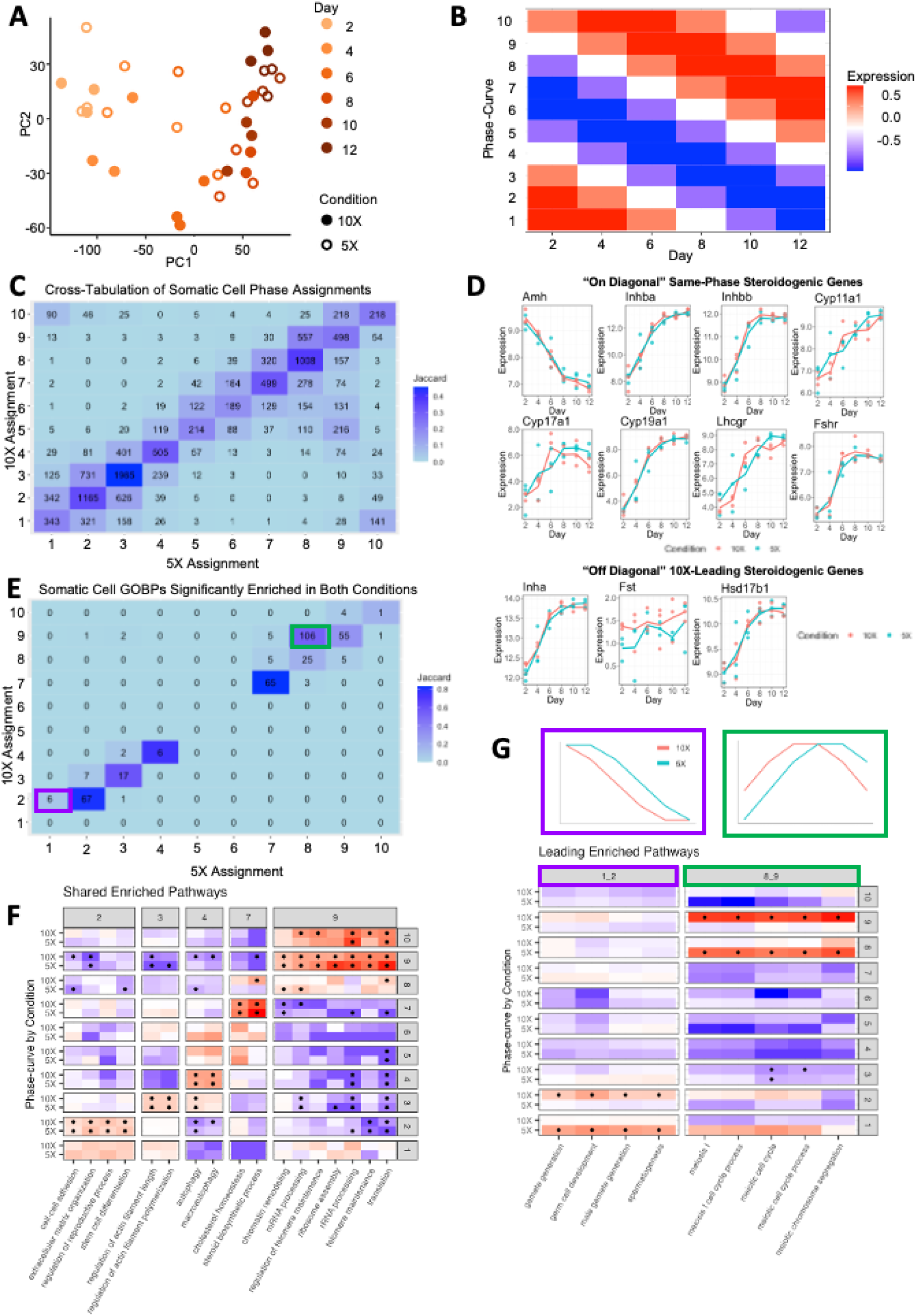
Profiling of the follicular somatic cell transcriptome over time *in vitro*. **A**) Principal component plot of the 36 somatic cell RNA samples, **B**) Expression of the 10 Phase-curves across the six timepoints, extracted from the sine function with steps of pi/5 and a length of 2*pi, **C)** Cross-tabulation of the Phase-curve assignment between 5X and 10X in somatic cells. Counts indicate the number of genes in the Phase pair. The pairs are colored by Jaccard Index (intersection over union) to show the similarity of the assignment between 5X and 10X (darker color indicates more similarity between the corresponding Phases), **D)** Expression of canonical steroidogenic genes across culture, with same-phase genes shown above and 10X-leading genes shown below, **E**) Same outlay as C, but instead of showing genes we show GOBPs assigned to the Phases. Highlighted tiles match 10X-leading pairs discussed in F, **F**) Highlighted pathways that have matched Phases between 5X and 10X. The x-axis are the highlighted pathways, split by the Phase which they are assigned (column groups). On the y-axis we plot the 10 Phases (row groups) split by 5X and 10X. Plotted in the heatmap are the log2 odds-ratios of those pathways for the Phases (values were limited to a range of (-2,2) to highlight contrasts between Phases). Stars within the tile indicate the odds ratio had an associated FDR < 0.1. Above the column groups the Phase-curves are plotted showing the change in expression over time, and **G**) Similar to F, but highlighting the corresponding (color boxes) 10X-leading pairs from E.

We used gene-clustering to identify the major temporal patterns exhibited by the DET genes. We calculated the six per-day expression values (averaged over the 3 replicates) for 5X, concatenated them with the six values for 10X, and clustered the genes using three approaches: SOM clustering with a 3x3 layout, K-means clustering with K=6, and supervised assignment to the family of 10 standard curves (see **Methods**), ultimately proceeding with supervised assignment of 13,580 genes to a family of 10 predefined standard curves representing gene expression patterns over time (Fig. 2B). For example, Phase Curves 1 and 2 can both be described as following a “high to low” trajectory of gene expression, yet Phase 2 starts to decrease earlier than Phase 1 (i.e., on Day 4 rather than Day 6). Similarly, Phase Curves 6 and 7 follow a “low to high” pattern, with Phase 7 increasing earlier than Phase 6 (Fig. 2B). Formally, the ten curves, each with six points, were extracted from a sine function, which completes a full 2*π cycle with steps of π/5, such that Curve-11 would have been the same as Curve-1. We retained Curves 1-10 as the complete reference set for the 10-way phase assignment.

We classified the 13,580 genes into one of the 10 phase-defined gene “clusters”, once by their 5X gene patterns and, independently, by their 10X gene patterns. After obtaining two sets of phase assignments for each gene, we examined phase-concordant or phase-discordant patterns between 5X and 10X by cross-tabulation of the two assignments (Fig. 2C). Nearly half (49%) of the DET genes were found along the diagonal in the phase tabulation, indicating they followed the same phase curve in 5X and 10X (Fig. 2C). Therefore, these genes reflect a common temporal signature of somatic cells that contributes to follicle growth in both conditions. Notable among this shared pool of genes are those known for driving important facets of folliculogenesis: *Amh* decreased over culture (Phase 1), while *Inhba, Inhbb, Cyp11a1, Cyp17a1, Cyp19a1, Lhcgr,* and *Fshr* increased (Phases 7 and 8), following expected patterns of steroidogenesis across development (Fig. 2D).

Aside from the same-phase genes, many genes were phase-advanced, or delayed, in one of the conditions. For some of these “off-diagonal” genes, the temporal expression change occurred earlier in 10X (i.e., the 10X-Leading category), while for others the change occurred earlier in 5X (10X-Lagging category). For example, the expression of steroidogenesis-related genes *Inha, Fst,* and *Hsd17b1* (which code for inhibin subunit alpha, follistatin, and hydroxysteroid 17-beta dehydrogenase-1, respectively) increased in both conditions, but were 10X-Leading: assigned to Phase 8 in 10X but Phase 7 in 5X (Fig. 2D and Supplemental File 1). Steroidogenesis is an important facet of follicle maturation, and the delay in activation of steroidogenic genes in 5X may be related to the suboptimal outcomes observed in 5X, including slower follicle growth, decreased hormone production, and immature oocytes after *in vitro* maturation^11^.

To augment the gene-level analysis, we performed enrichment analysis of biological process gene ontology (GO-BP) terms for each phase curve’s gene list for 5X and 10X separately (20 gene lists in all) (Fig. 2E and Supplemental File 4). This led to 20 sets of BP enrichment results (Supplemental File 3), where we focused on 1,563 GO-BP terms with greater than 50 and fewer than 500 genes. Since all GO terms have enrichment scores for each of the 20 gene lists, we assigned each GO term to one of the ten phases by its lowest enrichment FDR, for 5X and 10X separately. In all, 384 pathways satisfied FDR<0.1 and odds ratio (OR) >1 (to focus on enrichment rather than depletion) in at least one of the ten lists for both 5X and 10X and received phase assignments for both conditions. Using the same approach for gene-level phase comparison described above (Fig. 2C), we obtained cross-tabulation of 5X and 10X phase for GO-BP terms (Supplemental File 4) and found that the majority of the GO-related pathways have the same phase assignment (n=236) (Fig. 2E). Among these on-diagonal pathways are those related to steroid metabolism, steroid biosynthesis, and cholesterol metabolism, which were assigned to Phase 7 (Fig. 2F), showing increasing function over time in both 5X and 10X. Conversely, GO terms for "ECM organization" and "stem cell differentiation" (Phase 2 in Fig. 2F) and those related to "regulation of actin filament" (Phase 3) were enriched for Phase-2 and -3 genes, indicating that these functions decreased over time.

We then examined the pathways showing 10X-Leading or 10X-Lagging patterns (Fig. 2G), and found that most are 10X-Leading, i.e., above the diagonal in the phase tabulation. Terms related to "gamete generation" and "germ cell development" were assigned to Phase 1 in 5X and Phase 2 in 10X (left panel of Fig. 2G), representing genes with an earlier decline in 10X than 5X. Terms related to "meiotic cell cycle" were Phase 8 in 5X and Phase 9 in 10X (right panel of Fig. 2G), representing genes with earlier upregulation in 10X.

### Comparison with previously reported stage-specific granulosa and theca cells *in vivo* validates the dynamic expression of the somatic cells in the cultured follicles

A recent single-cell RNA-seq analysis of the adult mouse ovary by Morris et al.^12^ reported multiple granulosa and theca subtypes, which were annotated by their presence in follicles of different stages. As such, we were presented with an opportunity to compare the dynamic expression pattern for somatic cells in our follicle culture time series with the early- and late-stage granulosa and theca cells maturing *in vivo*, as annotated by Morris et al. First, we examined the dynamic pattern in our data for the top 10 marker genes reported by Morris et al. for the cell types most likely to match the primary to preovulatory stages of folliculogenesis recapitulated in our *in vitro* culture system. These included five of the granulosa cell subtypes (“Preantral-Cumulus”, “Antral-Mural”, “Mitotic”, “Atretic”, and “Luteinizing Mural” (Fig. 3A-E) and two of the theca cell subtypes (“Early Theca”, and “Steroidogenic Theca”) (Fig. 3F-G). We expected that the first list of markers, for "Preantral-Cumulus" granulosa cells, would follow a “high to low” pattern in our time series, as preantral granulosa cells differentiated into antral cumulus and mural cells^13,14^. Indeed, all nine genes (nine of the 10 were found in our study, *Gatm, Kctd14, Igfbp5, Col18a1, Pcsk6, Slc18a2 Fndc5, Wt1, Tmem184a*) decreased over time in our data (Fig. 3A). Conversely, all nine marker genes in the second list, for "Antral-Mural" (*Inha, Mro, Nap1l5, Hsd17b1, Slc26a7, Grem1, Cyp19a1, Tom1l1, Nppc*), increased over time (Fig. 3B), also consistent with the expectation that differentiated granulosa cells increase during culture. The third list, for “Mitotic” granulosa cells, also showed strong temporal specificity, as all ten genes (*Top2a, Ube2c, Racgap1, Birc5, Ccna2, Ccnb2, Hmgb2, Cdca8, Cdk1, Prc1*) uniformly peaked on Days 6-8 of culture (Fig. 3C). Collectively, this agreement between the two approaches: Morris et al.’s cell type-specific markers from scRNAseq analysis and our time-series data, offered reciprocal validation, reinforcing the reliability and robustness of both datasets.

**Figure 3:**
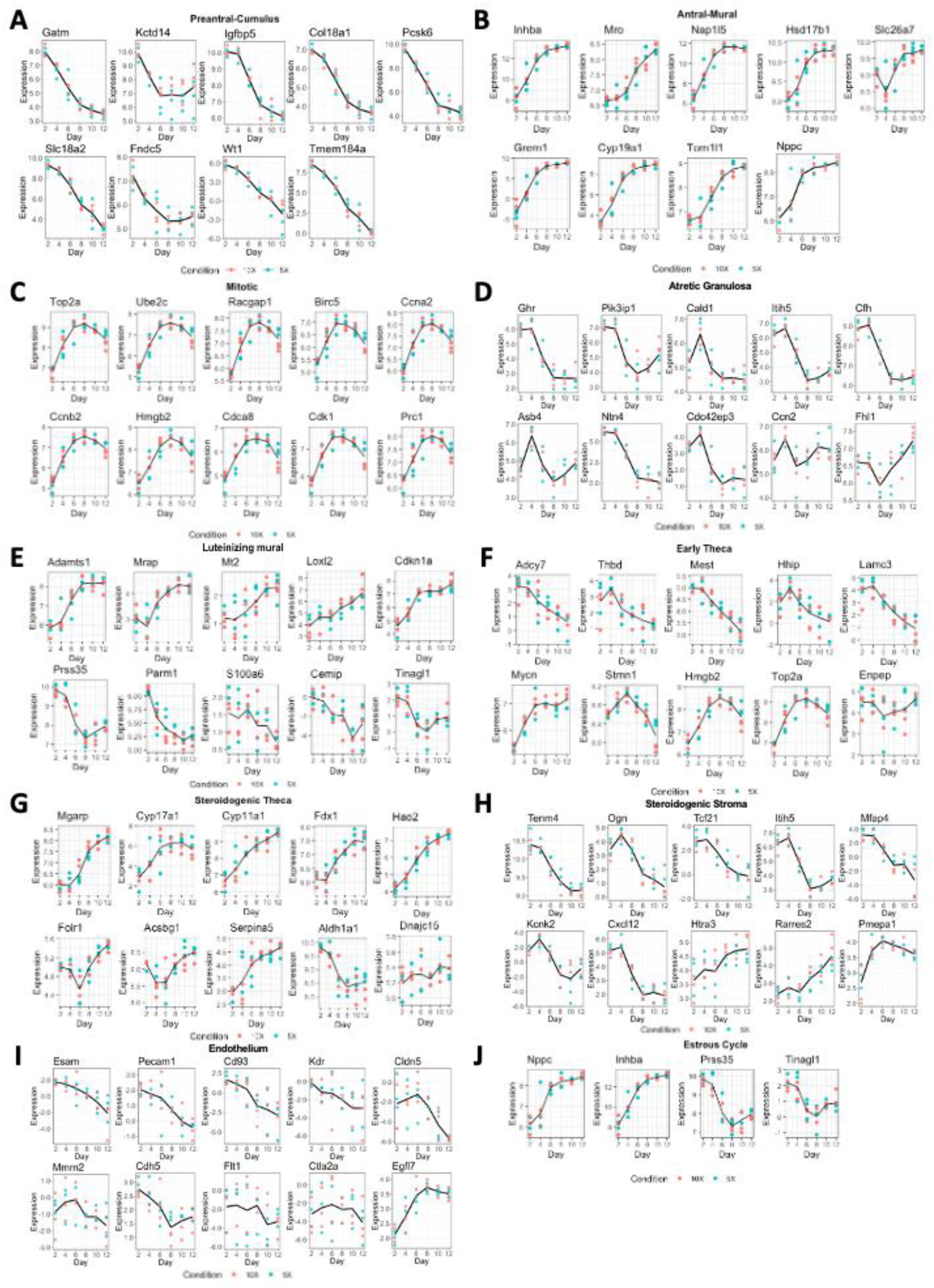
General concordance between the top 10 marker genes from *in vivo* single cell transcriptomics in mice (Morris et al.)^12^ and our *in vitro* follicles. **A**) Markers for “Preantral-Cumulus” granulosa cells with the expectation of decreasing expression over time (9/9 agreement). **B**) Markers for “Antral-Mural” granulosa cells with the expectation of increasing expression over time (8/8 agreement). **C**) Markers for “Mitotic” granulosa cells with the expectation of increasing then decreasing expression over time (10/10 agreement). **D)** Markers for “Atretic” granulosa cells showing 8/10 decreasing over time. **E**) Markers for “Luteinizing mural” granulosa cells with the expectation of increasing expression (5/10 agreement, 2/10 disagreement, 3/10 non-specific). **F**) Markers for “Early Theca” cells with the expectation of decreasing over time (5/10 agreement, 4/10 disagreement, 1/10 non-specific). **G**) Markers for “Steroidogenic Theca” with the expectation of increasing expressing over time (8/10 agreement, 1/10 disagreement, 1/10 non-specific). **H)** Markers for “Steroidogenic Stroma” with 7/10 decreasing over time and 3/10 increasing. **I)** Markers for “Endothelium” with 8/10 decreasing over time, 1/10 remaining constant, and 1/10 increasing. **J**) Markers for estrous cycle with two patterns (decreasing, increasing) appearing in our data.

The remaining two granulosa cell subtypes (“Atretic” and “Luteinizing Mural”) showed complex temporal dynamics in our culture. In the fourth list, for “Atretic” granulosa cells, eight of the ten genes (*Ghr, Pik3ip1, Cald1, Itih5, Cfh, Asb4, Ntn4, Cdc42ep3*) increased to their highest point on Day 4, followed by a decrease afterwards; while the other two genes (*Ctgf*/*Ccn2, Fhl1)* dipped after Day 4, yet increased in the last two time points between Days 10-12 (Fig. 3D). For the fifth and final list, "Luteinizing Mural" granulosa cells, five of the genes (*Adamts1, Mrap, Mt2, Loxl2, Cdkn1a*) showed increased expression (Fig. 3E), while the other five decreased (*Prss35*, *Parm1, S100a6, Cemip, Tinagl1*), although *S100a6* appeared noisy, and was not among the ANOVA-identified temporally-dynamic genes. The divergence may be explained by two unresolved subtypes of Luteinizing Mural granulosa cells, which possess distinct markers, or by the fact that our culture was not intended to drive the complete differentiation of these cells. We did not add human chorionic gonadotropin (hCG) or luteinizing hormone (LH) to our culture to induce oocyte maturation, and this may be one of the reasons that mural granulosa cells in culture did not express the full set of luteinization markers.

For the two lists of theca cell markers, five of the ten genes for "Early Theca" (*Adcy7, Thbd, Mest, Hhip Lamc3*) decreased over time (Fig. 3F), as would be expected for early theca precursor cells. However, one of the 10 (*Mycn*) showed increased expression, and three (*Stmn1, Hmgb2, Top2a*) increased at first, peaking in the middle days, and decreased again. The 10th gene (*Enpep*) was noisy and largely constant. For "Steroidogenic Theca" cells, eight of the ten genes (*Mgarp, Cyp17a1, Cyp11a1, Fdx1, Hao2, Folr1, Acsbg1, Serpina5*) increased over time, albeit with varying latency (Fig. 3G), as expected for these cells’ late appearance in development, with one gene (*Aldh1a1*) decreasing and one (*Dnajc15*) being noisy and uninformative. The increase of the eight steroidogenic theca markers is consistent with previous work from our group that reported androstenedione production and downstream synthesis of estradiol and progesterone, increased over twelve days of culture^10^.

We also examined the dynamic patterns for Morris et al.’s markers for “Steroidogenic Stroma” cells and “Endothelium”. Seven of the ten “Steroidogenic Stroma” genes (*Tenm4, Ogn, Tcf21, Itih5, Mfap4, Kcnk2, Cxcl12*) decreased over time, while three (*Htra3, Rarres2, Pmepa1*) increased (Fig. 3H). Similarly, eight of the ten “Endothelium” genes (*Esam, Pecam1, Cd93, Kdr, Cldn5, Mmrn2, Cdh5, Flt1*) decreased over time, while one (*Ctla2a*) remained mostly constant and one (*Egfl7*) increased (Fig. 3I). Finally, Morris et al. highlighted six genes for their distinct patterns across the Proestrus, Estrus, Metestrus, and Diestrus stages of the cycle, with four found in our data (*Nppc, Inhba, Prss35, Tinagl1*)^12^. Two of them, *Nppc* and *Inhba*, were reported in their study (Fig.6 of Morris et al.) as peaking in the Proestrus stage and decreasing afterwards, and they showed an increasing pattern in our data (Fig. 3J). Conversely, *Prss35* and *Tinagl1* peaked transiently in the Estrus stage in that study, and they had a high-low-high pattern in our data, reaching the lowest point in the middle stages of culture (Fig. 3J).

Overall, the observed concordance between the two studies is remarkable, especially considering that Morris et al. analyzed whole adult ovaries with follicles from all stages, whereas our data measured the progression of folliculogenesis from primary to preovulatory stages in culture over time. The temporal patterns for their marker genes in our data strongly supported their cell type annotation and the reported marker genes, and at the same time provided independent confirmation of the fidelity of our culture system.

### Dynamics of oocyte transcriptomes during culture

Parallel analysis of the oocyte samples, similar to the somatic cells (Fig. 2A), showed progressive transcriptional change from D2 to D12, and relatively small differences between 5X and 10X (Fig. 4A). Two-way ANOVA led to 10,091 DET genes (FDR<0.05), 14 DEC genes, and none for the Interaction term. For gene clustering of the 10,091 Day-varying genes, we replicated adopted analogous approaches as for the somatic cells, ultimately adopting the supervised assignment to the family of 10 standard curves.

**Figure 4:**
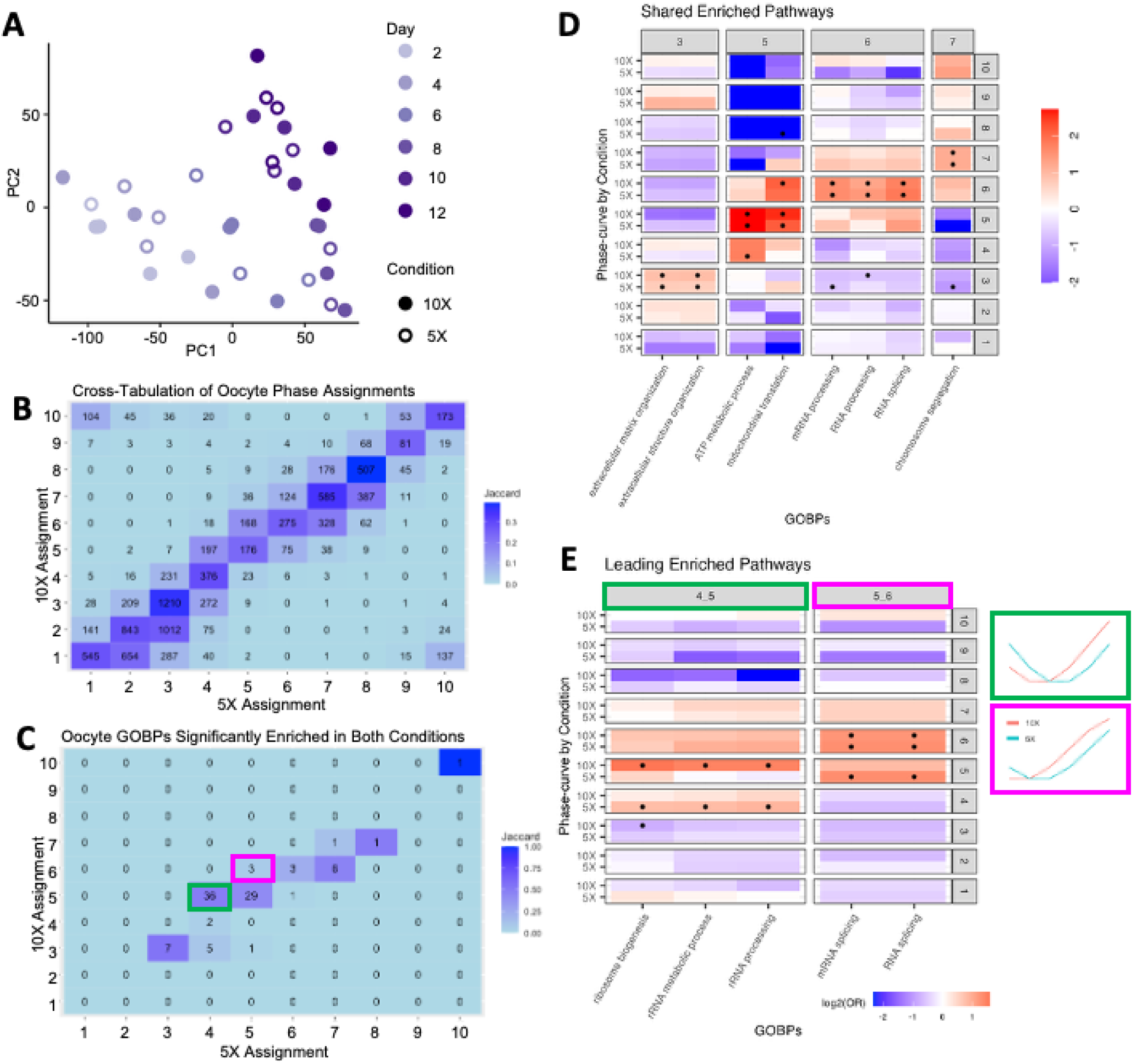
Profiling of the oocyte transcriptome over time *in vitro*. **A)** Principal component plot of the 36 oocyte RNA samples. **B**) Cross-tabulation of the Phase-curve assignment between 5X and 10X in oocytes. Counts indicate the number of genes in the Phase pair. The pairs are colored by Jaccard Index (intersection over union) to show the similarity of the assignment between 5X and 10X (darker color indicates more similarity between the corresponding Phases). **C**) Same outlay as B, but instead of showing genes we show GOBPs assigned to the Phases. Highlighted tiles match 10X-leading pairs discussed in E, **D**) Highlighted pathways that have matched Phases between 5X and 10X. The x-axis are the highlighted pathways, split by the Phase which they are assigned (column groups). On the y-axis we plot the 10 Phases (row groups) split by 5X and 10X. Plotted in the heatmap are the log2 odds-ratios of those pathways for the Phases (values were limited to a range of (-2,2) to highlight contrasts between Phases). Stars within the tile indicate the odds ratio had an associated FDR < 0.1. Above the column groups the Phase-curves are plotted showing the change in expression over time, and **E**) Similar to D, but highlighting the corresponding (color boxes) 10X-leading pairs from C.

After we assigned the 10,091 genes to the same set of 10 standard curves described above (Fig. 2B), for 5X and 10X data separately, we proceeded to compare the genes’ 5X and 10X phase assignments. Similar to somatic cells, almost half of the dynamic genes in oocytes (47%) were “on the diagonal” (n=4,771), having the same phase in 5X and 10X, while 44% (n=4,423) were shifted by one, “off the diagonal” (Fig. 4B).

As above, we performed GO-BP enrichment analysis for the 20 lists of genes, representing the 10,091 genes’ assignments to 10 phases based on their patterns in 5X and 10X, separately. Few terms were significantly (FDR<0.1) enriched (OR>1) in at least one of the ten gene sets in both 5X and 10X conditions (n=96, Supplemental File 4). These included 43 same-phase pathways, 39 10X-Leading pathways, and 14 10X-Lagging pathways (Fig. 4C). The same-phase pathways included those related to “extracellular matrix organization” (Phase 3), “ATP metabolic process” and “mitochondrial translation” (Phase 5), “RNA processing” and “RNA splicing” (Phase 6), and “chromosome segregation” (Phase 7) (Fig. 4D).

All 39 10X-Leading pathways were enriched for genes with late activation (Phases 4, 5, and 6), but with earlier activation in 10X. Specifically, GO terms related to ribosome function were enriched for Phase-4 genes in 5X and Phase-5 genes in 10X (left panel, Fig. 4E). Similarly, GO terms related to mRNA splicing increase earlier in 10X (Phase 6) than 5X (Phase 5) (right panel, Fig. 4E). Accumulation of ribosomes has been linked to oocyte developmental competence^15,16^. Therefore, a delayed accumulation of mRNAs and ribosomes in the oocytes cultured in 5X may contribute to their reduced ability to mature at the end of culture, and future work may target ways to increase ribosome biogenesis in the 5X condition and validate its correlation with maturation outcomes.

### Condition-significant genes in somatic cells and oocytes

As described above, two-way ANOVA for somatic cell data found 306 genes with an overall significant difference between 5X and 10X conditions. To investigate the temporal pattern of these DEC genes, we calculated the six per-day expression values (averaged over the 3 replicates) for 5X, concatenated with the six values for 10X, and clustered the 306 genes by their 12-point series. SOM clustering (**Methods**) with a 2-by-2 layout yielded four clusters (Fig. S1C), with genes in each cluster submitted to *PANTHER*^17^ for GO enrichment analysis (**Methods**). An analogous approach was used to analyze the 728 Interaction-significant genes (Fig. S1D). DEC genes in two of the four clusters were expressed higher in 10X over all six time points (Fig. S1C), including those with roles in the reproductive process (*Mroh2b, Brdt, Umodl1, Sebox, Wee2, Xlr3a, Bmp15*) (Supplemental File 1), and those related to exocytosis and vesicle-mediated signaling *(Syt1, Cspg5*) also had higher expression in 10X. Genes in the other two clusters were either consistently lower in 5X or "switched" in the middle of the time course.

Compared to somatic cells, far fewer genes were significant between 10X and 5X conditions in oocytes (n=14). Of these, five have biological annotation (*Hexdc, Slc10a3, Unc45a, Zfp383, and Olfr624)* and were expressed higher in 10X than 5X (Fig. S1E), with only *Hexdc* having a known function in the oocyte; it encodes hexosaminidase-D, one of a family of proteins important for sperm-egg interaction during fertilization^18^. Of note, *Olfr624,* an olfactory receptor without reported roles in the ovary, was overall higher in 10X in both the somatic cell and oocyte data (Fig. S1E, shown again in S1F for individual samples, before averaging the 3 technical replicates). *Olfr624* is among the several olfactory receptors showing differences in somatic cells between 5X and 10X cultured primary murine follicles in a previous microarray-based study from our lab^11^. Olfactory receptors have been suggested by other studies for potential involvement in sperm chemotaxis during fertilization^19–21^ and could possibly have similar roles in ovarian follicles. Overall, the greater number of condition-significant genes in somatic cells (306 vs. 14 in oocytes), suggests that two conditions had a stronger impact on somatic cells during follicle growth and maturation.

### Dynamics of oocyte-somatic cell interaction by analyzing ligand-receptor pairs

Bidirectional communication between the oocyte and the somatic cells is essential for coordinated development of the follicle. In particular, early stages of folliculogenesis are driven by paracrine signaling, as demonstrated by group culture studies^9–11^. Since we collected matched somatic cell and oocyte datasets over multiple timepoints, we sought to investigate temporal changes in mediators of paracrine signaling by analyzing joint expression levels of ligand-receptor (L-R) pairs. We relied on an online repository for L-R pairs in mice^22^, which matched our RNA-seq data to 647 ligand genes and 343 receptor genes, for 2,422 L-R pairs (as some ligands and receptors are represented multiple times in the 2,412 pairs). We organized L-R analyses by four categories (Fig. 5A): "Cell-Cell" - both the ligand and the receptor expression data in somatic cell data, "Oocyte-Oocyte" - both in oocytes, "Oocyte-Cell" - pairing the ligand in oocyte and its receptor in somatic cells, and "Cell-Oocyte" - the reverse pairing.

**Figure 5:**
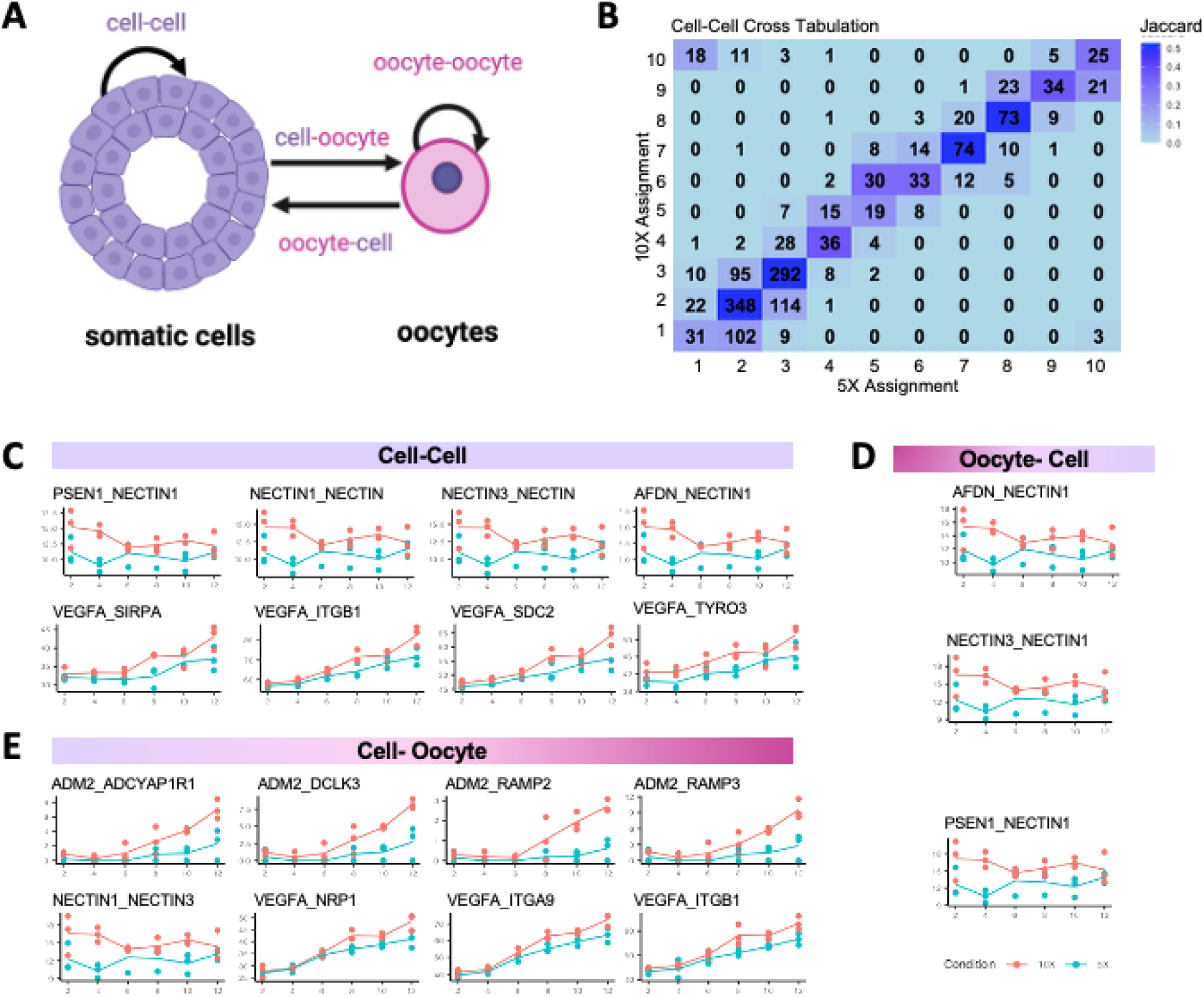
Ligand and receptor pairs between and within somatic cells and oocytes. **A**) Schematic of the four communication types: Cell-Cell, Oocyte-Oocyte, Oocyte-Cell, and Cell-Oocyte, **B**) Crosstabulation of assigned Phase-curves between 5X and 10X for all pairs in the Cell-Cell category. Boxes indicate groups of 10X-leading and 10X-lagging pairs. Crosstabulations for all other categories shown in Supplemental Figure S2, and **C**) Scatter and line plot of the expression across days of selected Condition-significant L-R pairs in Cell-Cell, **D)** Oocyte-Cell, and **E)** Cell-Oocyte. Plot titles include the communication type and name of the L-R pair. Figure 5A was generated using Biorender.com.

**Figure 6:**
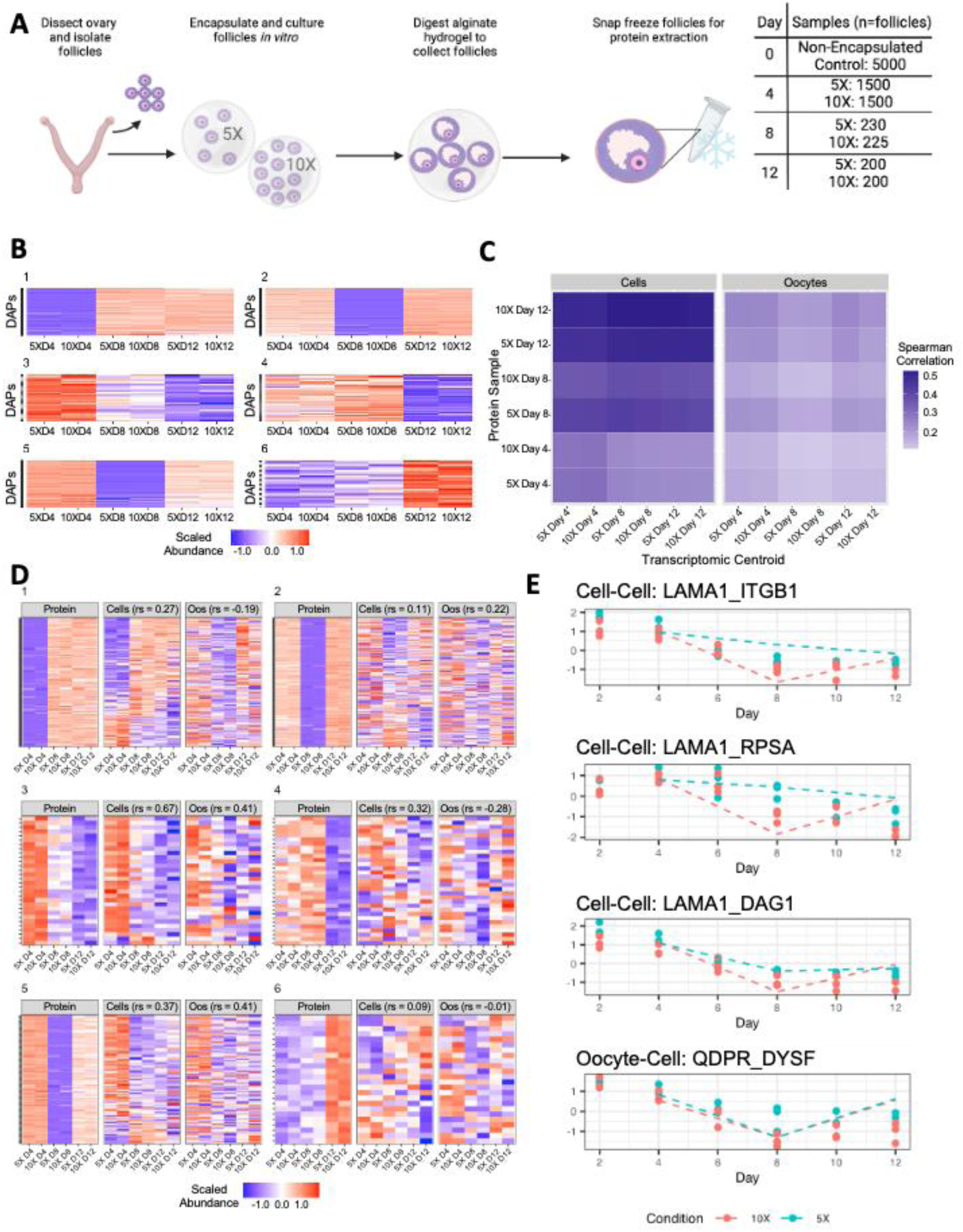
Proteomic profiling of follicles over time *in vitro*. **A)** Schematic describing the experimental design, beginning with primary follicles cultured in alginate hydrogels in groups of 5 (5X) or 10 (10X). To collect samples for proteomics, hydrogels were enzymatically digested, and the entire follicle was snap frozen for analysis, **B**) Six clusters of differentially abundant proteins across Days. In order, they show the following patterns across three timepoints consistent between 5X and 10X: low-high-high, mid-low-high, high-mid-low, mid-high-low, high-low-mid, and low-mid-high, **C**) Heatmap showing the correlation of matched centroids in the RNA data to the samples from proteomics. Spearman correlation on overlapping features colors tiles, **D**) Clusters of DAPs by day plotted alongside expression from paired centroids from transcriptomic data (somatic cells and oocytes). Proteins/gene pairs are ordered by decreasing correlation between protein abundance and gene expression in cells, from top to bottom. Average spearman correlation is shown in the headers for cells and oocytes within each cluster, **E**) The four DEIs from transcriptomics that are present in the protein ligand and receptor analysis, colored by condition. Points represent scaled interaction expression in the indicated interaction, with lines indicating the expression of the proteomic data. Figure 6A was generated using Biorender.com.

For each of the four categories and for each of the 2,422 L-R pairs, we calculated a “communication score” as the product of the log expression levels of the gene pair (**Methods**). This was done for the six time points (Days 2-12), two conditions (5X and 10X), and three replicates (Table S1), thus for each of the four categories we obtained a 36-by-2,422 data matrix, which served as a new sample-feature data table containing 2,422 L-R features and 36 samples of the same design. By following the same strategy for analyzing somatic cell and oocyte gene expression levels, we ran two-way ANOVA on the L-R scores, with Day, Condition, and an Interaction term as factors (results in Supplemental File 5). Some of the DE L-R pairs were significant in more than one category (e.g., in both Cell-Cell and Cell-Oocyte) (Fig. S2A).

For the temporally significant L-R pairs, we followed the same procedure as above to assign the dynamic phase of an L-R pair, then compared its phase between 5X and 10X by cross-tabulation (Cell-Cell cross-tabulation shown in Fig. 5B, and the other three categories in Fig. S2B-D). Most L-R pairs had concordant phase (on-diagonal), while some others showed leading or lagging relationships. For example, among the Cell-Cell temporally significant L-R pairs, a majority (n = 965, 61%) showed the same phase in 5X and 10X, with 288 as 10X-leading, and 324 as 10X-lagging (Fig. 5B). These L-R results, organized in Supplemental File 5, offer a deeper view than the phase analyses for individual genes (Supplemental File 1). A pair of ligand-in-cell and receptor-in-oocyte, for instance, might peak in the middle of the 10 phases *and* show 10X-leading, i.e., peak earlier in 10X condition than in 5X. For example, the somatic cell ligand *Igf1* and oocyte receptor *Igf1r* pair followed the Phase-2 trajectory in 5X (high to low) and Phase-9 (mid-high-mid) in 10X. Meanwhile, the somatic cell ligand *Igf1* and oocyte receptor *Igf2r* pair followed the same Phase-10 (increasing followed by decreasing) trajectory in both conditions. *Igf1* has important roles in oocyte maturation,^23^ and the diverging relationships between *Igf1* and its receptors in 5X and 10X uncovered in the L-R analysis provided a more nuanced understanding of somatic cell-oocyte signaling than the individual gene analysis. Examining this crosstalk in the other direction, oocyte ligands *Gdf9* and *Bmp15* did not appear in the oocyte DET genes, but they were represented in the significant oocyte-cell L-R pairs. *Gdf9*-*Orai2* and *Bmp15*-*Bmpr1a* followed 10X-lagging trajectories (5X Phase 2 and 10X Phase 1) while *Bmp15*-*Bmpr1b* followed a 10X-leading trajectory (5X Phase 7 and 10X Phase 8). *Gdf9* and *Bmp15* are both oocyte-secreted factors that encourage granulosa cell proliferation and theca cell differentiation as a follicle develops.^24^

Examining the differences between 5X and 10X, 32 L-R pairs were significant across conditions (Supplemental File 5), including four pairs with ligand *Adm2* in the Cell-Oocyte category (Fig. 5E), seven pairs with ligand *Vegfa* in the Cell-Cell and Cell-Oocyte categories (Fig. 5C,E), and sixteen cell-adhesion L-R pairs with *Nectin1, Nectin3, Afdn,* and *Psen1* in several categories (Fig. 5C-E). *ADM2* ligand pairs showed higher levels in later days of culture, likely driven by rising expression of ADM2 itself. Pro-angiogenic factor *Vegfa* consistently showed increased expression with time in culture that rose higher in 10X. Cell adhesion-related L-R pairs showed consistently higher expression in 10X.

### Intrafollicular proteomic profiling of follicles across culture timepoints

To augment the RNA-seq analysis, we performed shotgun proteomics for 5X and 10X cultured follicles on Days 4, 8 and 12, as well as a D0 baseline sample, for a total of seven samples (Fig. 6A). The reduced sampling in these experiments was due to the large amount of material needed to perform proteomic analyses (**Methods**). In all, 5,066 proteins were detected (with nonzero abundance values) in at least one of the seven samples, with 3,096 proteins being abundant at multiple timepoints (in either 5X, 10X, or both).

Since proteins varied greatly in the number of samples in which they were detected, we separated the analysis in two parts. First, for the 1,970 proteins detected only in one of the three time points (left side of Fig. S3A), we submitted those seen in D4, D8, or D12 as three separate lists for GO analysis to identify enrichment patterns in the early, middle, or late stages of murine folliculogenesis. Second, for the remaining 3,096 seen in multiple time points (right side, Fig. S3A), we performed two-way ANOVA to identify proteins with significant temporal changes or significant differences between 5X and 10X. We did not include an Interaction term, as there is no replicate sample in the 2-by-3 design.

For the proteins only detected at one timepoint, the D4, D8, and D12 lists contained 32, 4, and 1,002 proteins, respectively. Their GO-BP enrichment results are in shown in Supplemental File 6. In transcriptome data, *Amh* RNA was expressed early in culture, and its protein appeared only in Day 4. The D12-only proteins were depleted for genes involved in mitochondrial organization and ATP synthesis and enriched in cilium organization (Supplemental File 6). The most notable finding was the enrichment for cilium-related function in D12 proteins, as 41 of the 71 genes annotated for “cilium organization” appeared among the 1,002 D12-only proteins. It has been previously reported that granulosa cells in antral follicles possess cilia, and knockdown of cilium function negatively impacts ovulation^25^.

For proteins detected in multiple time points, ANOVA identified 468 proteins significant for temporal changes (FDR<0.1), and none across conditions (Supplemental File 7). We then clustered the 468 differentially abundant proteins (DAPs) for Day using K-means (K=6), leading to six clusters containing 25 to 135 genes (far right column in Fig. S3A, with their expression heatmaps in Fig. 6B). Cluster-1, with 135 proteins, followed a low-high-high pattern and included FST (follistatin), INHBA and INHBB (inhibin β-A and inhibin β-B) (Fig. S3B), which are members of the TGFβ super family and regulate granulosa cell proliferation and differentiation starting at the preantral stage^26^. Cluster 3 (n=30) contained proteins with a decreasing pattern, and included ZP2 and ZP3 (zona pellucida glycoproteins 2 and 3) (Fig. S3B), consistent with a previous report of widespread presence of ZP1/2/3 in human primordial follicles^27^. Temporal changes of these ZP proteins over the course of follicle development have not been previously reported, and they showed similar trends in our RNA dataset (Supplemental File 1). Cluster 4 (n=36) contained proteins with higher abundance on D4 and D8, and a drop on D12. While this list did not contain any proteins with known links to folliculogenesis, it contained those associated with mitochondrial function and ATP synthesis: NDUFA2, NDUFA12, NDUFB4, NDUFS1, COA3, and SCO2 (subunits of NADH dehydrogenase [ubiquinone] 1 alpha and beta subcomplexes, NADH-ubiquinone oxidoreductase core subunit 1, cytochrome c oxidase assembly factor 3, and synthesis of cytochrome c oxidase 2) (Fig. S3B). This is consistent with the result in the single-timepoint analysis, showing D12-specific proteins are depleted for those for mitochondrial organization and ATP synthesis. Due to the reduced number of timepoints (from 6 to 3), we did not run phase assignment with the proteomic data.

To investigate the relationship between mRNA and protein patterns, we first mapped the 3,096 multi-time detected proteins to the genes in transcriptome data, finding 2,924 in both. Since the protein data had only 3 time points and no replicates, we extracted the transcriptomic data for the corresponding two time points and three replicates, averaged them to obtain the centroids for the three time points matching protein samples, and for the two conditions. This reduction of RNAseq data was done for cells and oocytes separately, and each would be compared with protein data, which did not separate somatic cells from the oocytes.

We ran three sets of protein-RNA comparisons. First, to assess the protein-RNA relationship at the sample level, we calculated the Pearson’s correlation coefficients (using the 2,924 genes) between each of the 6 protein centroids and each of the 12 RNA centroids, where six were for somatic cells and six were for oocytes (Fig. 6C). Proteomic centroids showed a higher correlation with the somatic cell RNA centroids than with the oocyte RNA centroids, generally showing a higher correlation at later timepoints (Fig 6C). This result was unsurprising, since the protein analysis used whole-follicle samples, which were dominated by the material from the somatic cells, especially at later timepoints, when the follicles grow larger and have even higher representations of somatic cells.

Second, to compare at the gene level, we used proteins with significant dynamic patterns and juxtaposed the protein heatmaps (left of the three panels in Fig. 6D) with those for RNA in Cells (middle panel) and RNA in Oocytes (right panel), for genes in each of the six protein clusters shown in Fig. 6B, using the same gene order over the set of three heatmaps. As in the sample-level comparisons (Fig. 6C), protein data for individual genes showed a greater correspondence with the somatic cell transcriptome than with the oocyte transcriptome (Fig. 6D). Even for the RNA data in somatic cells, the similarity with protein data varied: strong for many genes in Cluster-1 and weak in other genes and in other clusters.

Third, we compared protein and RNA in L-R analysis. There were 190 L-R pairs in the protein data, involving 72 ligands and 66 receptors. Since proteomic samples were from pooled follicles, not separating somatic cells and oocytes, the data only allowed the calculation of one category: Follicle-Follicle L-R “communication scores”, for the three timepoints and two conditions. ANOVA of the 190 L-R features and six samples found 59 pairs DET proteins (FDR<0.1), and none for Condition. We then focused on the 32 DEC L-R pairs from the transcriptomic data. Of these, four were present in the proteomic data, and all were higher in 5X, with mostly consistent dynamic patterns between RNA and protein (Fig. 6E, with dotted lines showing protein data). The results for these four L-R pairs provided mutual validation between RNA and protein.

## DISCUSSION

In this study, we performed unbiased transcriptomic and proteomic analyses of the murine 5X/10X group follicle culture system, which was developed as a fully controlled, reproducible model for studying folliculogenesis *in vitro*^5–7^. Our goal was to characterize the molecular changes along a twelve-day, seven-point time course, with paired samples for oocytes and their surrounding somatic cells, and to compare between 5X and 10X group cultures.

To assess the progression of cultured follicles *in vitro* relative to the known stages of naturally matured follicles *in vivo*, we leveraged multiple gene lists generated recently by Morris et al.^12^ Their group investigated the transcriptome of ovaries in cycling adult mice, which contained follicles at different developmental stages, including early-stage and antral stage follicles and postovulatory corpora lutea^12^. The data presented here serves as a strong complement to those by Morris et al., as we profiled a carefully size-selected population of primary follicles undergoing folliculogenesis in a coordinated fashion for 12 days *in vitro*. Between the two datasets, we observed an overall striking concordance of the temporal profile for most of the marker gene sets generated by Morris et al., including five gene sets of the granulosa cell clusters (“Preantral-Cumulus”, “Antral-Mural”, “Mitotic”, “Atretic”, and “Luteinizing Mural”) and two gene sets of the theca cell clusters (“Early Theca” and “Steroidogenic Theca”) (Fig. 3). This result provided mutual validation to both studies. For Morris et al., which analyzed ovaries collected at distinct timepoints during the estrous cycle, our results independently confirm the gene activity dynamics in a true time-series dataset. For our study, the concordance in Fig. 3 strongly suggests that in our cultured follicles, granulosa cells can indeed differentiate into more mature stages as seen *in vivo*. Thus, our culture system, by faithfully recapitulating the temporal progression of follicles maturing in vivo, stands as a valid experimental platform for meticulously studying both the inter- and intracellular interactions that drive folliculogenesis, and serves as a promising method for translational fertility preservation in the future.

A key advantage of the follicle culture system is the ability to examine the temporal dynamics of folliculogenesis from the primary to pre-ovulatory stages. While previous studies have reported on the transcriptional signature of murine ovaries *in vivo*^12,28^; oocytes from primordial, transitioning primordial, and primary follicles^29^, and cumulus-oocyte complexes across *in vitro* maturation^30^, each of these provides a cross-sectional view of follicle development, rather than a time course across development. To our knowledge, this is the first attempt to collect paired-sample time series for oocytes and somatic cells from group cultured follicles, offering an unparalleled opportunity to study leading- and lagging-temporal relationships between these two cellular compartments. Notably, our controlled culture system ensures that the data we present is unique to follicles encapsulated all at the primary starting stage, without confounding factors from other cell types or stages of follicles.

Our main analysis strategy involved two-way ANOVA for Condition (5X and 10X) and Time (the six time points from D2 to D12) and revealed strong dynamic changes for both the oocytes (10,091 genes) and follicular somatic cells (13,461 genes). In comparison, the genes with significant changes between 5X and 10X were much fewer (14 genes for oocytes and 306 genes for somatic cells), and many of them were challenging to interpret due to Interaction effect and the impact of batch correction (see Methods). We analyzed the dynamic changes at both the gene and pathway levels, with many results aligning with past knowledge (Fig. 2-4). While highlighting some of the most notable genes and pathways for both somatic cells and oocytes, we also collated the full set of data and results and shared them as a community resource (Supplemental Files 1-4).

Somatic cells in both 5X and 10X showed the same decreasing expression of *Amh* over culture (Supplemental File 1). *Amh* is a member of the TGFβ family that suppresses primordial follicle activation, regulates follicle sensitization to follicle stimulating hormone (FSH), and suppresses *Cyp19a1* production for estradiol synthesis^31–33^. *Amh* is typically secreted by the granulosa cells in preantral and early antral follicles followed by a decrease in large antral follicles. In contrast, we saw an increase in steroidogenesis-related genes *Inhba, Inhbb, Inha, Fst*, *Cyp11a1, Cyp17a1, Cyp19a1, Hsd17b1, Lhcgr,* and *Fshr* (Fig. 2D). At the pathway level, somatic cells of both 5X and 10X showed an increase in gene sets related to steroid metabolism, steroid biosynthesis, and cholesterol metabolism (Fig. 2F), which indicates that *in vitro* grown follicles ramp up steroid hormone production as culture progresses. We also found a decreasing pattern for “extracellular matrix organization” (Fig. 2F). The extracellular matrix plays a vital role in folliculogenesis, participating in processes including primordial follicle activation, maintenance and support of follicle structure, retention of secreted paracrine factors, and steroidogenesis^34–37^. ECM organization decreased at the pathway level, reflecting a highly dynamic extracellular environment, which has also been reported by Grosbois et al. Their recent study showed that the ECM of human ovarian tissue changed over six days of *in vitro* culture, including a significant decrease in collagen^37^. This data can be applied for development of biomimetic materials that can interact with the changing ECM during follicle culture^38,39^.

Within the somatic cell temporal analysis, we also investigated the changing expression of markers for theca and endothelial cells. In our study, follicles were isolated at the primary stage, with the oocyte surrounded by one layer of granulosa cells enclosed by the follicle’s basement membrane. Thus, we retained an insignificant or possibly completely absent number of theca and endothelial cells, which are present only outside the basement membrane^40^. Nevertheless, the concentration of androstenedione that reached approximately 0.4 ng/mL in the culture media by Day 12 in our previous studies^10^ and the expression of “Early Theca” and “Steroidogenic Theca” cell markers in this study (Fig. 3F-G) suggest the presence of theca-like cells. We propose two explanations. First, it is possible that a few theca cells were retained during the isolation process, and if that is the case, it is clear that the current follicle culture platform is capable of supporting theca cell survival, proliferation, and differentiation throughout culture. Second, our alternative explanation is that granulosa cells within the follicle may have the capacity to act as bi-potential cells, differentiating into theca cells or acting as theca-like by secreting androstenedione. Granulosa cells of enzymatically digested murine follicles have previously been shown to express *Cyp17a1*, a known theca-specific cell marker, by the end of culture, providing some support for this theory^41^. Steroidogenic theca-like cells expressing *Cyp17a1* have also been described as “Steroidogenic Stroma” by Morris et al.^12^. Future work should validate the spatiotemporal expression of theca markers in group culture of primary follicles through *in situ* hybridization or immunofluorescent staining to better characterize whether cells are adopting bi-potential characteristics.

In contrast to the theca, expression of endothelial cell marker genes decreased over time (Fig. 3I), suggesting that the current design of the follicle culture system is unable to support endothelial survival (Supplemental File 1). Vascular endothelial cells play a crucial role in follicle development *in vivo*, responding to paracrine cues from the follicles and intercalating into the theca layer. *In vitro*, ovarian endothelial cells have been shown to support follicle survival and growth^42^. Therefore, future work should investigate new methods for incorporating endothelial cells or endothelial cell-secreted factors into the group follicle culture platform.

Though there were much fewer genes with significant differences between 5X and 10X Conditions as described above (Supplemental File 1), we highlight a few results that may account for the better outcomes in the 10X cultures. Follicles in the 5X and 10X conditions achieved similar rates of survival (Fig. 1C), growth over time (Fig. 1E), and appeared morphologically normal (Fig. 1F), so we expected that the transcriptional profile of these conditions would contain a high percentage of overlap. Our results agree with our previous microarray data^11^, indicating that a relatively small, yet critical number of differentially expressed genes accounted for the superior outcomes in 10X. The transcriptome comparison between 5X and 10X somatic cells identified reproductive processes-related genes (*Mroh2b, Brdt, Umodl1, Sebox, Wee2, Xlr3a, Bmp15*), which had greater expression in 10X than 5X throughout culture (Supplemental File 1). *Sebox*, skin-, embryo-, brain-, and oocyte-specific homeobox, is expressed in the oocytes of growing, non-atretic follicles starting at the primary stage and peaking in antral stages^43^. Similarly, Wee1-like protein kinase 2 (*Wee2,* also known as *Wee1b)* is found in oocytes and increases throughout folliculogenesis. *Wee2* plays a role in maintaining meiotic arrest in mice^44,45^ and rhesus macaques^46^, and mutations in the *Wee2* gene have been linked to infertility in humans^47,48^. Finally, bone morphogenetic protein 15 (*Bmp15)* is an important paracrine signaling factor secreted by oocytes to promote granulosa cell proliferation during folliculogenesis^49^. Though these are canonical oocyte markers, *Bmp15* has previously been reported in the follicular fluid and granulosa cells of follicles collected for *in vitro* fertilization, with decreased levels associated with age and poor response to stimulation^50^. Together, these 10X-associated genes may partially explain the superior follicle growth and maturation observed in 10X. The somatic cell analysis also revealed new, additional genes and pathways, including expression of genes involved in vesicle-mediated signaling (e.g. *Syt1, Cspg5*). Vesicle-mediated signaling has recently received attention in the study of folliculogenesis, especially given the importance of signaling via secreted factors in the follicular fluid of antral follicles^51,52^. In fact, one group reported that *Syt1* is involved in multiple stages of oocyte development, including the meiosis I to meiosis II transition and the calcium-mediated cortical granule reaction during fertilization^53,54^. Somatic cell expression of this gene in our dataset may have a related role but has yet to be investigated. Furthermore, there were only 14 differentially expressed genes across conditions in our oocyte data. This suggests that the phenotypic differences observed *in vitro*, i.e., the differential oocyte maturation outcomes between 5X and 10X^11^ could be driven by the somatic cell compartment of the follicle, not the oocyte. Thus, future efforts to develop a translational follicle culture system may need to be particularly focused on supporting proper somatic cell development, which in turn supports oogenesis. Experiments targeting the 14 DEC genes in oocytes reported here may show whether somatic cell support drives follicle outcomes independent of oocyte transcription, or if oocyte transcriptome changes are somehow directing somatic cell gene expression.

We identified 32 L-R pairs with significant differences across conditions, including repeated expression of ligand *Adm2 (*Fig. 5E). A recent study has implicated the role of *Adm2* in folliculogenesis by regulating the synthesis of estradiol^55^. Ligand *Vegfa* also appeared several times (Fig. 5C,E), confirming the crucial role *Vegfa* plays in ovarian follicle development. *Vegfa* has been previously reported as a differentiating factor between 5X and 10X follicles^9,10^. Furthermore, half of the differential 5X vs. 10X L-R pairs were related to cell adhesion, including pairs with *Nectin 1, Nectin 3, Afdn,* and *Psen1 (*Fig. 5C-E). Nectins have significant effects on male fertility and spermatogenesis^56^ and have previously been localized in the ovary^57–59^, possibly suggesting a role in normal folliculogenesis. Since these findings are based on co-expression of the receptor and the ligand in RNA data, they remain to be validated using quantitative proteomic analysis. It is also important to note that the transcriptome-based L-R analysis does not encompass many non-protein signaling molecules, such as lipids and steroid hormones, which also require further investigation in the future.

We expected to observe some concordance between our proteomic and transcriptomic datasets; however, we also expected to see discrepancies, as not all mRNAs are transcribed to proteins, and there may be temporal delays between transcription and sufficient accumulation for detection at the protein level. Notably, our proteomic data validated gene expression patterns identified in our transcriptome data for the key regulators of folliculogenesis FST and INHBA/INBHH, and zona pellucida glycoproteins, ZP2 and ZP3 (Fig. S3B). These biological processes and individual proteins may be used as targets for exogenous modulation to support follicle development *in vitro*. Future iterations of this work may focus on the secretome by profiling the spent media of follicles cultured *in vitro*, but the challenge of removing serum from culture media remains to be solved, as it is an essential factor for follicle development^60^.

In summary, this study contributed novel datasets for mouse follicles cultured *in vitro* - paired oocyte and somatic cell transcriptome data over a 12-day time series, and shotgun proteomics data. Such data have been lacking in the field yet are crucially needed to provide a more detailed understanding of the dynamics of follicle developmental progression. We analyzed temporally dynamic and differentially expressed genes, pathways, and proteins, and assessed L-R expression as a way to profile the direction and temporal change of putative cellular interactions. The analysis results and all the raw and derived data are shared, and they have augmented our prior analysis of the system based on microarrays. We found that more genes were significantly differentially expressed over time, rather than between conditions, and that somatic cells may drive differences across conditions and thus the differential follicle maturation outcomes observed in culture. Future work building off this study should focus on understanding targeted mechanisms of paracrine communication during follicle development, ultimately advancing *in vitro* follicle growth and maturation as a translational fertility preservation method.

## METHODS

### Follicle Isolation, Encapsulation, and Culture

Primary ovarian follicles were isolated, encapsulated in alginate matrices, and cultured as previously described^10,11^. Briefly, whole ovaries were removed from C57B6 X CBA/J mice ages 10-12 days. All animal procedures were performed in compliance with the Guidelines for the Care and Use of animals at the University of Michigan and approved by the Animal Care and Use Committee at the University of Michigan (PRO00009635). Ovaries were washed in warm Leibovitz’s L-15 medium (L-15) (Thermo Fisher) with 0.5% (v/v) PenStrep (Thermo Fisher) and transferred to a dish with pre-equilibrated alpha modification of minimum essential medium (αMEM) (Thermo Fisher) with 0.5% (v/v) PenStrep. To collect large numbers of primary follicles from the ovaries, 10% (v/v) Liberase DH at 13 Wünsch units/mL (Sigma) was added to the dish of ovaries and gently mixed, then incubated undisturbed at 37°C for 35-45 minutes. After incubation, the dishes were pipetted for 5 minutes to break up the enzymatically-digested tissue, releasing individual follicles, then the digest was arrested using 10% (v/v) fetal bovine serum (FBS) (Thermo Fisher).

Primary follicles with diameters ranging 90-110 µm were selected from the ovary digest and encapsulated in 10 µL alginate beads in groups of 5 (5X) or 10 (10X) follicles per bead. To crosslink alginate, beads were dropped into a solution of 50 mM CaCl_2_ and 140 mM NaCl and allowed to crosslink for 3 minutes. Encapsulated follicles were cultured in 96 well plates with 150 µL growth media (GM) per well, containing αMEM supplemented with 3 mg/mL bovine serum albumin (BSA) (MPBiomedicals), 1 mg/mL bovine fetuin, 5 µg/mL insulin, 5 µg/mL transferrin, 5 ng/mL selenium (ITS, Sigma), and 10 mIU/mL highly purified, human-derived follicle-stimulating hormone (FSH) (Urofollitropin, Ferring Pharmaceuticals). Every 48 hours after encapsulation, half of the GM (75 µL) was replaced with fresh GM. Follicle growth was tracked by imaging follicles every 48 hours before exchanging GM. Follicle diameter from each 48-hour time point was measured using ImageJ by taking the average of two perpendicular measurements across the follicle. Hydrogels without exactly 5 or 10 fully encapsulated follicles were removed from culture.

Hydrogels containing follicles that fell out of the alginate hydrogel onto the bottom of the well, or hydrogels with follicles dead at the start of the culture (on day of encapsulation) were excluded from analysis and sample collection for sequencing. Follicles were termed “dead” if the oocyte was extruded more than 50% from the follicular structure with no surrounding granulosa cells, or if the follicular structure was dark and the oocyte was not visible. Only follicles surviving to Day 12 of culture were included for growth analysis, while all follicles were included in survival analysis. The results presented were pooled from 4 separate culture experiments, with a total of 2,920 follicles contributing to RNA samples for sequencing.

For growth analysis (Fig. 1E), follicles surviving to Day 12 of culture (n = 160 follicles from 10X gels, n = 140 follicles from 5X gels) were evaluated using a two-way ANOVA with repeated measures followed by a Tukey test to correct for multiple comparisons in Graphpad Prism 10. The same follicle data for follicles cultured up to Day 12 (n = 160 for 10X, n= 140 for 5X) was processed in Graphpad Prism 10 using the log-rank test for comparison of survival curves (Fig. 1D).

### Follicle Dissection, RNA Extraction, and Sequencing

Follicles (n = 910 for 5X, n = 1,110 for 10X) were taken from culture for dissection into somatic cell and gamete counterparts at the following time points: Days 2, 4, 6, 8, 10, and 12. Fresh, un-encapsulated follicles were also collected for sequencing as Day 0 controls, totaling nine sets of 100 follicles pooled from 20 mice. The number of follicles isolated for each RNA sample are outlined in Supplemental Table 1, with an average of 43 follicles/sample for 5X, 53 follicles/sample for 10X, and 100 follicles/sample for controls. To dissect follicles, alginate beads were digested by incubating with 10 IU/mL alginate lyase for 20 minutes. Follicles were then transferred to warm L-15 and oocytes were mechanically removed from the follicles using insulin syringes (BD 305620) then rinsed 3 times in L-15 to remove any remaining somatic cells before collection in a microcentrifuge tube. The remaining somatic cells were collected into a microcentrifuge tube. Oocyte and somatic cell samples were centrifuged at 300g for 5 minutes, supernatant was removed, and cell pellets were snap frozen in liquid nitrogen then stored at - 80°C. RNA extraction on each sample was performed using the Qiagen RNEasy Micro Kit following the manufacturer’s instructions. Extracted RNA was stored at -80°C before being submitted to the University of Michigan’s Advanced Genomics Core for sequencing. Samples were sequenced on the Illumina NovaSeq platform at a sequencing depth of 25 million reads per sample.

### RNAseq Data Analysis

The sequence data was analyzed in R Statistical Software (v 4.2.1)^61^. The initial count matrix contained 55,426 genes for 90 samples. We then reduced the number of genes using multiple filtering criteria. Firstly, retaining only genes that were present (non-zero counts) in at least half the samples in oocytes (n = 45) or in somatic cells (n = 45). We retained 25,348 genes from this step. After preliminary downstream analysis, we decided to also use a minimum standard deviation of raw expression of 10 and a minimum mean expression of 10 in either somatic cells or oocytes. The combination of all filters resulted in a dataset containing 16,927 genes. Next counts were normalized to counts per million for each sample and then log2 was taken with values floored at 0.02 (chosen because it was half the lowest CPM value of ∼0.04).

The data was viewed jointly with both cell types (cells and oocytes) by calculating the principal components (PCs) on the full log-transformed CPM matrix. The data was then viewed separately for each cell type. For further analysis we removed the day zero control since they were extreme outliers as the day zero follicles were more reflective of *in vivo* grown follicles and had not undergone the encapsulation and culture process. Batch effects were noticed in PC plots when viewing experimental batches in each day set (2-4, 6-8, 10-12). To adjust for this, we took the centroids for each batch (see Supplemental Table 1 for batch information). Then within each batch we subtracted the batch centroid from the four samples, generating batch corrected data. Batch effects were no longer present in PC projections and models (not shown). We then moved forward with the batched corrects PCs and models.

For the models, the linear model was calculated for each gene with Day as a categorical variable, Condition (5X vs 10X) and the Interaction of time and condition. ANOVA was then run on the results of the linear model to determine differentially expressed genes (DEGs) (Benjamini-Hochberg adjusted p-value < 0.05) for Day, Condition and the Interaction. For somatic cells DEGs we had 15,183 for Day, 306 for Condition, and 728 for Interaction. For oocytes DEGs we had 10,091 for Day, 14 for Condition, and zero for interaction. Results are in Supplemental File 1.

Kohonen’s self-organizing maps (SOM) (R package som) were used to group the differentially expressed genes. The average expression for all DEGs was calculated for each Day and Condition combination (n = 12). Maps of dimensions of 2 by 2 were used for Condition- and Interaction-significant DEGs. Condition-significant somatic cell DEGs had cluster occupancies of 55 (0_0), 128 (0_1), 89 (1_0), and 34 (1_1). Interaction-specific somatic cell DEGs had cluster occupancies of 191 (0_0), 47 (0_1), 40 (1_0), and 450 (1_1).

### Phase-shift Analysis

First, by using Self-Organization Map (SOM) with a 3-by-3 layout we divided the 13,461 genes into nine clusters (Fig. S1A, with cluster assignments in Supplemental File 1). The centroids of the nine SOM clusters were combined into four major groups, for patterns of gene expression that were temporally low to high (low-high), low-high-low, high-low, and high-low-high. In a slightly revised SOM approach, we obtained the nine SOM-based gene clusters for the 5X and 10X data separately, each using their 6-point series (rather than concatenating into a 12-point series). This allowed us to examine the genes’ concordant and discordant assignments between 5X and 10X in terms of high-low, low-high dynamic patterns (results not shown). In a further refined SOM approach, we concatenated the 5X and 10X data series "vertically", to build a dataset with each gene appearing twice, one for 5X and one for 10X, such that the SOM clustering of the 13,461*2 = 26,922 genes assigned the 5X gene patterns and 10X gene patterns into the nine clusters in a single SOM analysis, to achieve better 5X-10X comparability than running SOM separately (results not shown). We omit detailed description of the results here, as we progressed to two other approaches, described below, and found them to be more powerful.

In the second approach, we performed K-means clustering (K=6) to divide the 13,580 genes (increased from 13,461 DET genes to 13,580 when adding the 728 Interaction-significant genes (DEI)) into six clusters, for 5X and 10X separately, using the corresponding six-point temporal series described above (Days 2, 4, 6, 8, 10, and 12 of culture). The 10X centroids are shown in Fig. S2B, with cluster assignments for both 5X and 10X in Supplemental File 1. The resulting assignments into the six K-means clusters were consistent with the aforementioned nine SOM clusters (not shown), and largely consistent between 5X and 10X. Notably, the six cluster centroids (Fig. S1B) appeared to be "advancing" sequentially: from low-high in Cluster 1, to low-high-low in Cluster 2, and so on, to high-low-high in Cluster 6.

This observation, from unsupervised K-means clustering of day-varying genes, suggested that the major dynamic patterns can be categorized as a series of sequentially connected, phase-defined principal curves. If we classify the 5X gene patterns and, independently the 10X gene patterns, into a standardized family of phase-defined gene sets we would be able to identify (1) early, middle, or late-changing genes and (2) phase-concordant and phase discordant genes between 5X and 10X. This led to the next approach, which relies on supervised assignment of the 13,580 DET+DEI genes by their match to a family of 10 standard curves, each containing six equally spaced time points (Fig. 2B). The curves are ordered from 1 to 10, with a constant phase shift between the adjacent pairs. Formally, the ten curves, each with six points, were extracted from a sine function, which completes a full 2*pi cycle with steps of pi/5, such that Curve-1 is the same as Curve-11, and we retained Curves 1-10 as the reference set for the 10-way phase assignment.

We assigned each gene’s phase, 1-to-10, by its highest Pearson’s correlation coefficient among the 10 curves. The assignment is robust, as the second highest correlation always correspond to an adjacent curve for all the genes (i.e., never to a curve two or more shifts away). The 10-way assignment results for both 5X and 10X are in Supplemental File 1 and showed concordance patterns with the SOM’s 9-way assignment and the K-means 6-way assignment (not shown, but can be verified by cluster IDs in Supplemental File 1). The advantage of the supervised 10-way assignment, importantly, is that the 10 curves have known, precisely ordered phase shifts, while SOM and K-means clusters are not guaranteed to have precise are influenced by the relative density (i.e., number of genes) among clusters. Additionally, with the supervised assignment to 10 standard curves we can compare assignment across multiple datasets (cell transcriptomics, oocyte transcriptomics, L-R interactions).

### Pathway Analysis on RNA

Gene lists identified through SOM were submitted to the PANTHER database for gene ontology enrichment analysis^17^ with the background list of all 16,927 genes in the analysis. Genes were filtered through the “biological process” GO terms using gene name and tested using Fisher’s Exact t-test. Pathways were filtered to those that had > 50 and < 500 genes present in the background list. GO terms with FDR < 0.1were considered significant.

For gene lists identified through their assignment to the 10 phase-defined reference patterns for 5X and 10X, we submitted to PANTHER following the same protocol. Additionally, we focused on enriched rather than depleted pathways, thus adding the criterion of OR>1 to assigned pathway-phase. Pathways were further analyzed if they had an FDR <0.1 in at least one of the l0 lists for both 5X and 10X. For visualization in of pathway patterns in Fig. 2 and Fig. 4, odds ratios were limited to a range of (-2,2) to highlight contrasts between Phases.

### Ligand and Receptor Analysis

Ten ligand-receptor (L-R) datasets were downloaded from Github^22^. Datasets were harmonized and merged together. Unique L-R pairs with both the ligand and receptor present in our data were retained (n = 2,422). The pairs consisted of 647 ligands and 343 receptors. To calculate the communication score for a set of cells, we multiplied the batch-corrected expression of the ligand and receptor in the corresponding cells. Specifically, we either calculated the score between the matched somatic cells and the oocyte that were together before separation and sequencing (Cell-Oocyte and Oocyte-Cell), or we multiplied the ligand and receptor expression within each cell type/sample (Cell-Cell and Oocyte-Oocyte). For each time point, we then have three communication scores in 5X that pair uniquely to three 10X communication scores.

We next ran ANOVA for each L-R category (n = 4) across all L-R pairs (n = 2,422) on the linear model with covariates Day, Condition and Interaction. P-values were corrected using the Benjamini-Hochberg method, and significance was determined at FDR < 0.05. Temporal dynamics/phase shift analysis was then performed similar to previous sections. Each L-R category was then analyzed separately. We then broke down the cross-tabulation for pseudo-curve counts into three stages: early, mid, or late (Fig. 5B and S2B-D). This was determined by the expression pattern of the pseudo-curves in the time series data. To determine if a ligand or receptor was enriched for a stage, we performed a 3x2 Fisher’s exact test with the alternative hypothesis greater than and corrected the p-values to FDR. If a ligand or receptor was determined to be enriched (FDR < 0.10) at a stage, we then performed the same Fisher’s exact test but instead of testing stage we were testing if it was enriched in matched, 10X-leading, or 10X-lagging. All nominated ligands and receptors were each plotted on the cross-tabulation to confirm their enrichment in a time stage & matching/10X-leading/10X-lagging pattern (data not shown).

### Proteomics Sample Preparation and Profiling

Follicles were cultured in 5X and 10X as previously described for liquid chromatography tandem mass spectrometry (LC-MS/MS)-based proteomics. On Days 4, 8, and 12 of culture, follicles were isolated from alginate hydrogels as previously described, rinsed in DPBS-/-, and centrifuged at 300g for 5 minutes to form a cell pellet. Supernatant was removed and pellets were snap frozen and stored at -80°C. Fresh primary follicles (mice ages 10-12 days) were isolated and snap frozen as a control sample (n = 5,000 follicles). The number of follicles isolated for each protein sample, which were compiled over multiple culture experiments, are as follows: D0 control n = 5000, 5X D4 n = 1500, 10X D4 n = 1500, 5X D8 n = 230, 10X D8 n = 225, 5X D12 n = 200, 10X D12 n = 200. Each sample contained approximately 5 million cells based on predicted number of cells in follicles of various diameter. Cell pellets were delivered to the University of Michigan Proteomics and Peptide Synthesis Core for analysis. Protein extraction was performed in modified RIPA buffer (2% SDS, 150mM NaCl, 50mM Tris pH 8, 1X Roche complete protease inhibitor) using mechanical disruption with NextAdvance buller blender with 1mm stainless steel beads. The protein concentration of each sample was determined by Qubit fluorometry (Invitrogen). 20μg for each sample was processed by 5cm SDS-PAGE using a 10% Bis-Tris Novex mini-gel (Invitrogen) and the MOPS buffer system. The mobility region was excised into 20 equally sized bands and each was processed by in-gel digestion using a robot (DigestPro, CEM) with the following protocol: washing with 25mM ammonium bicarbonate followed by acetonitrile; reduction with 10mM dithiothreitol at 60°C followed by alkylation with 50mM iodoacetamide at RT; digestion with trypsin (Promega) at 37°C for 4 hours; quencing with formic acid. The supernatant was analyzed directly without further processing. Half of each digested sample was analyzed by nano LC-MS/MS with a Vanquish Neo HPLC system interfaced to a ThermoFisher Fusion Lumos mass spectrometer. Peptides were loaded on a trapping column and eluted over a 75μm analytical column at 350nL/min; both columns were packed with PepMap Neo C18 resin (ThermoFisher). The mass spectrometer was operated in data-dependent mode, with the Orbitrap operating at 60,000 FWHM and 15,000 FWHM for MS and MS/MS respectively. The instrument was run with a 3s cycle for MS and MS/MS. 10 hours of instrument time was used for the analysis of each sample.

### Proteomics Data Processing

Data were searched using a local copy of Mascot (Matrix Science) with the following parameters: Enzyme: Trypsin/P; Database: SwissProt Mouse (concatenated forward and reverse plus common contaminants); Fixed modification: Carbamidomethyl (C); Variable modifications: Oxidation (M), Acetyl (N-term), Pyro-Glu (N-term Q), Deamidation (N/Q); Mass values: Monoisotopic; Peptide Mass Tolerance: 10 ppm; Fragment Mass Tolerance: 0.02 Da; Max Missed Cleavages: 2. Mascot DAT files were parsed into Scaffold (Proteome Software) for validation, filtering and to create a non-redundant list per sample. Data were filtered using 1% protein and peptide FDR and requiring at least two unique peptides per protein, providing spectral counts for each sample.

Across all samples 5,066 proteins were identified as abundant in at least one of the seven samples and used as the initial abundance matrix for further data analysis. Day 4 and 8 samples showed high levels of missingness compared to Day 12 samples. Proteins that only had abundance at a single timepoint (ignoring Control samples since they were not included in downstream ANOVA testing) were filtered and analyzed separately. The remaining 3,086 proteins were then normalized by the total protein abundance for each sample then multiplied by a factor of 10^6^ before being floored at 1 and then log2 normalized. A two-way ANOVA test was then performed using Condition (5X or 10X) and Day as a categorical variable as the inputs. False discovery rates (FDRs) were then calculated from the p-values using the Benjamini-Hochberg correction. Differentially abundant proteins (DAPs) were determined by an FDR < 0.10, with 468 proteins passing this threshold for Day and none for Condition. The DAPs for Day were then clustering into 6 clusters using k-means clustering. In the proteomic output provided by Scaffold, conversions between proteins and genes were provided and utilized for matching protein and RNA samples.

For ligand and receptor analysis we performed a similar analysis as with the transcriptomic data. Because somatic cells and oocytes are not separated in the proteomic data, all pairs were calculated on follicle-to-follicle interactions. 190 pairs were identified in the proteomic data covering 72 ligands and 66 receptors. Of the 190, 173 were also present in the transcriptomic data for comparison. Running ANOVA on Day and Condition (we lacked degrees of freedom to look at Interaction), we found 50 differentially abundant for Day and none for Condition.

### Pathways Analysis on Proteins

Gene and protein lists were submitted to the PANTHER database for gene ontology enrichment analysis^17^. The reference list for the day-specific analysis the full list of 5,066 proteins were used, and for the cluster analysis the 3,086 proteins tested with ANOVA were used as the reference list. Genes were filtered through the “biological process” GO terms using gene symbol, and proteins were filtered using UNIPROT ID. Each gene/protein list was evaluated and tested using Fisher’s Exact t-test. GO terms were restricted to terms with at least 50 proteins and no more than 500 proteins. After restricting the number of terms for analysis, FDRs were calculated using Benjamini-Hochberg correction. GO Terms with FDR < 0.10 were considered significant.

## Supporting information

Supplemental File Readme

## ACKNOWLEDGMENTS

We thank the members of the Shikanov and Li laboratories for scientific discussions and manuscript feedback. We would also like to acknowledge the University of Michigan’s Advanced Genomics Core and the Proteomics & Peptide Synthesis Core for RNA and proteomics processing, respectively, and their staff for guidance and initial data analysis.

---

**Supplemental Figure 1:**
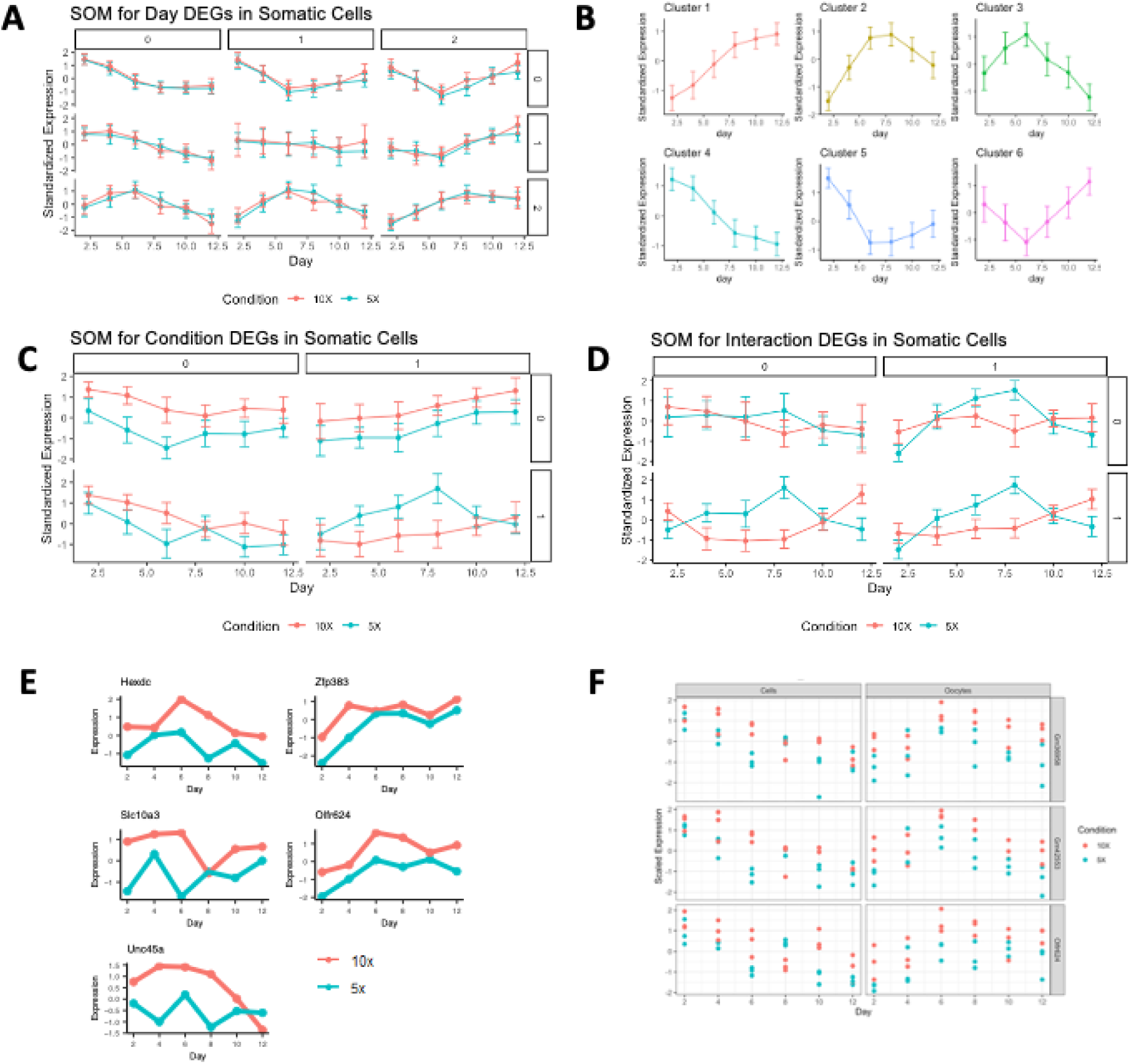
Profiling of the somatic cell and oocyte transcriptomes *in vitro*. **A**) Centroids of the nine SOM clusters of the 13,461 temporally significant genes in somatic cells. **B**) K-means (K = 6) clustering result of the 13,580 Day- or Interaction-significant genes in somatic cells, showing ordered "phase advance" between adjacently numbered clusters. **C)** Centroids of the four Self-Organizing Map (SOM) gene clusters, in a 2-by-2 arrangement, for the 306 Conditional-significant genes in somatic cells, showing both within-cluster mean and standard deviation, with color indicating 5X and 10X data. **D)** Centroids of the four SOM clusters, in 2-2 format, for the 708 Interaction-significant genes in somatic cells. **E)**. Average expression of the 5 Condition-significant DEGs in oocytes with biological annotation. All show greater expression in 10X compared to 5X. **F**) Scatterplot of the three genes that were Condition-significant DEGs in both somatic cells and oocytes, with concordant greater expression in 10X.

**Supplemental Figure 2:**
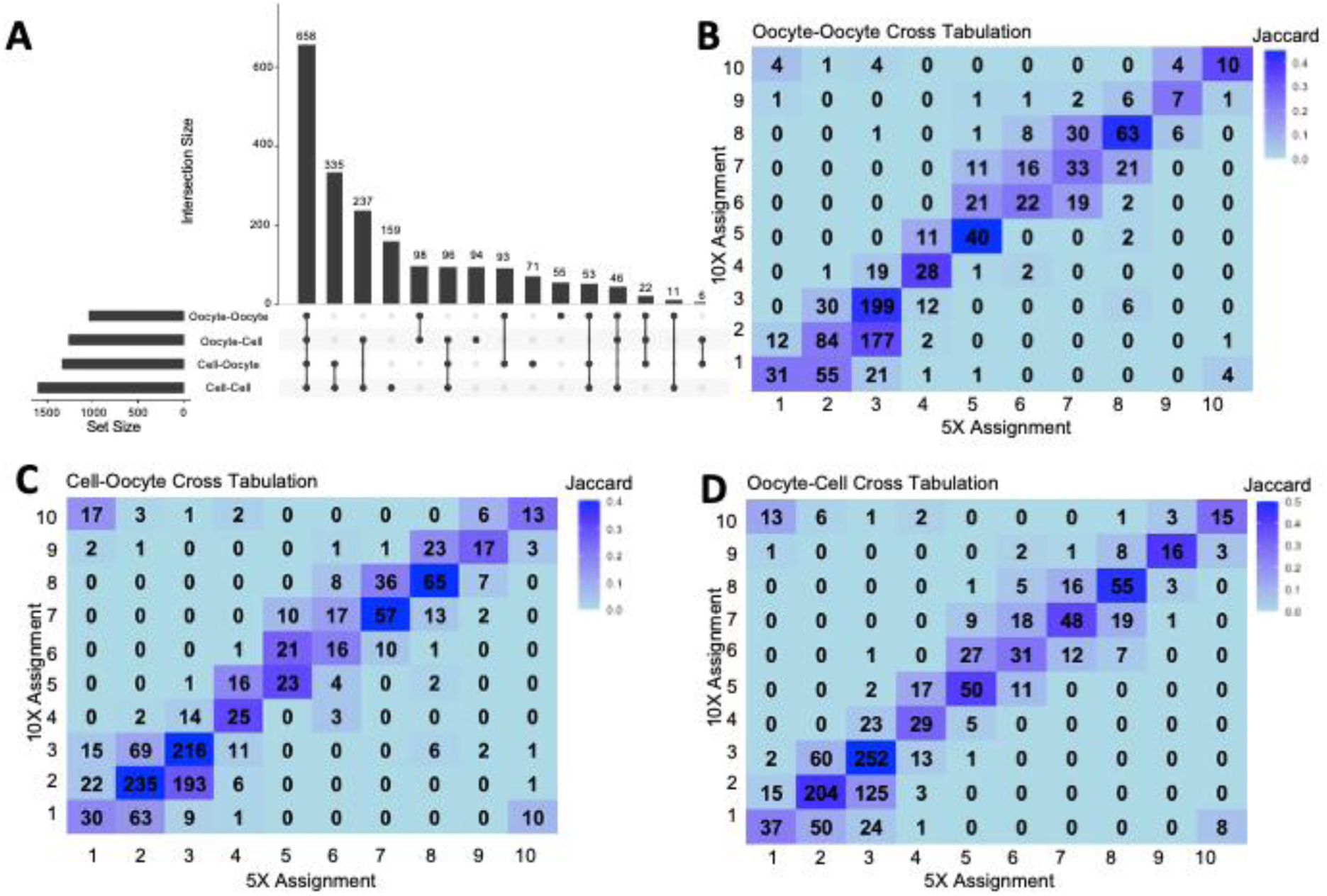
A) Upset plot of the Day-significant interactions. The bottom left bar chart shows the number of Day-significant DEGs for each of the four interaction types. The intersection size bar chart indicates the number of elements shared across the sets (indicated by the interaction chart below). For example, the first vertical intersection bar indicates there are 658 interaction pairs that are present in all four Day-significant interaction lists, **B**) Crosstabulation of assigned Phase-curves between 5X and 10X for all interactions in Oocyte-Oocyte, **C)** Cell-Oocyte, and **D)** Oocyte-Cell.

**Supplemental Figure 3:**
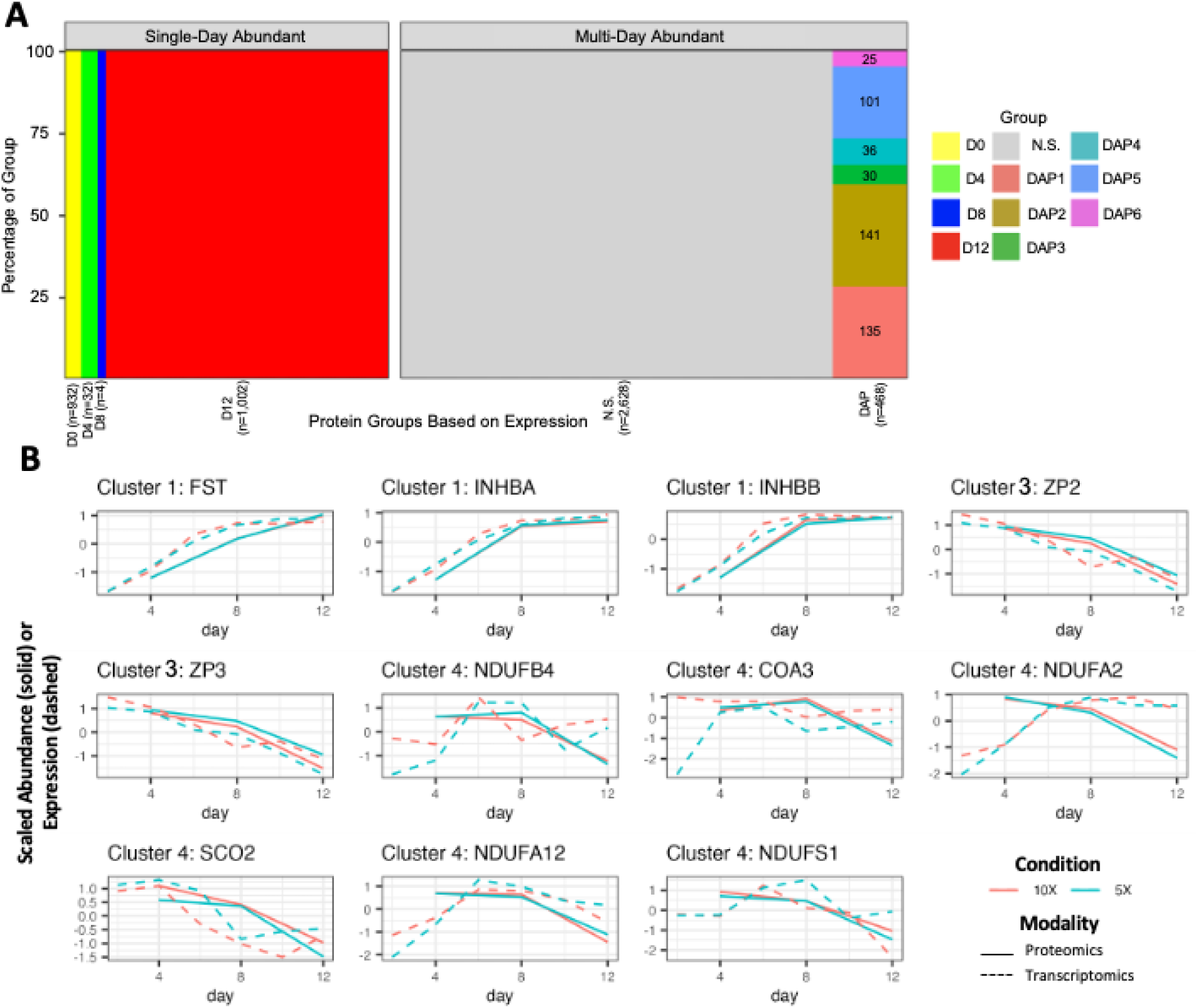
Proteomic profiling of follicles over time *in vitro,* cont. **A**) Diagram representing the distribution of proteins into different categories for analysis. The x-axis divides the 5,066 proteins into 100 bins, with the width groups equal to the percentage of the proteins it represents. The first four columns show proteins expressed at a singular timepoint. Multi:N.S., represents genes that were expressed at multiple timepoints but were not differentially expressed proteins (DAPs). Proteins that were DAPs were then divided into 6 clusters using K-means clustering. The number of proteins per category are listed in the column label on the x-axis. **B**) Line plots for the standardized abundance/expression by Day and Condition for the specific proteins/genes noted in analysis. Lines are color by Condition, with proteomics data in solid lines, and average transcriptomic data in dashed lines.

**Supplemental Table S1:**
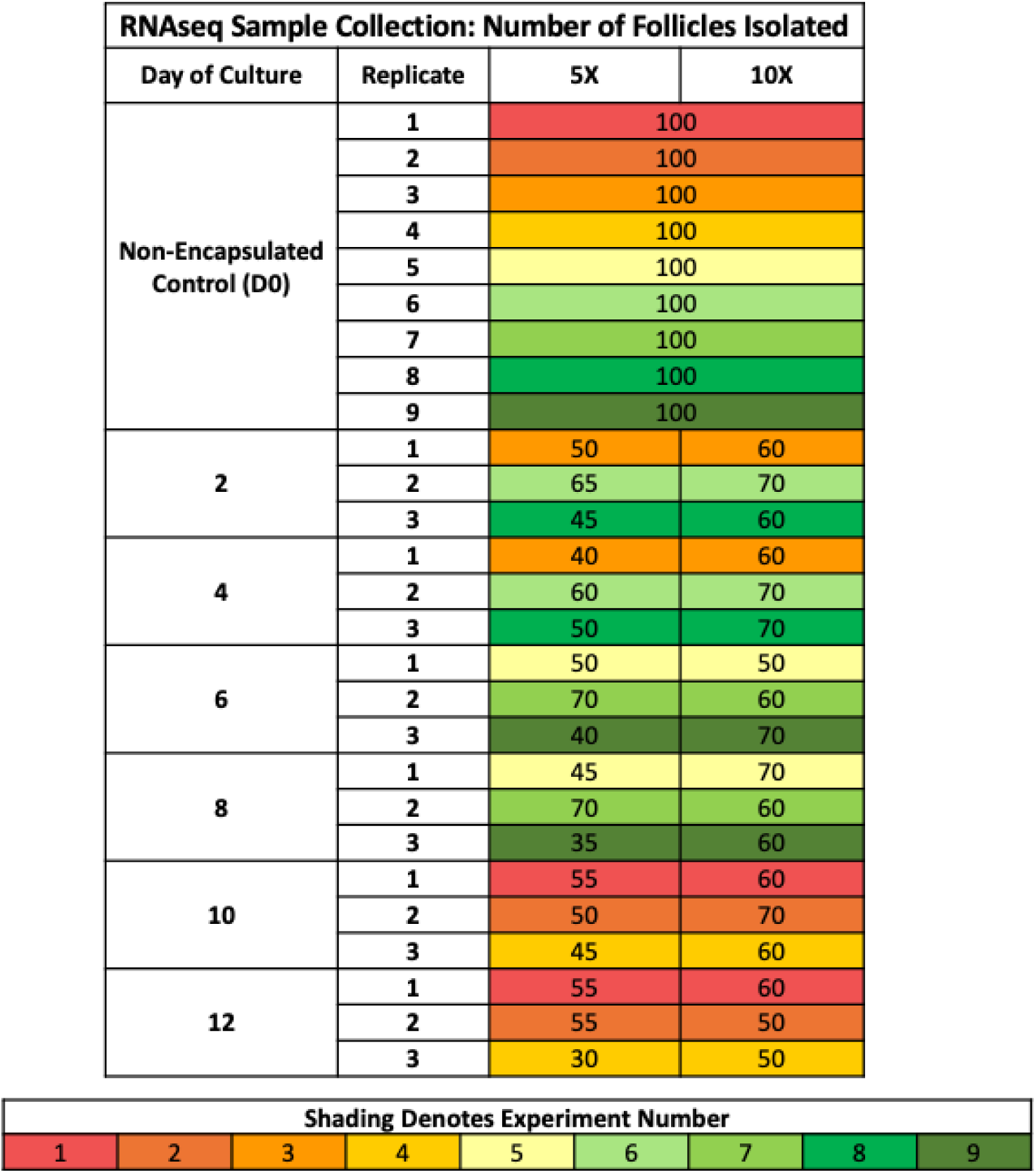
A schematic showing the number of follicles isolated for each day of culture across the three replicates (and 9 fresh controls). Shading represents that batch they were sequenced in, across the samples, conditions, and timepoints.

| RNAseq Sample Collection: Number of Follicles Isolated |  |  |  |
| --- | --- | --- | --- |
| Day of Culture | Replicate | 5X | 10X |
| Non-Encapsulated Control (D0) | 1 | 100 |  |
|  | 2 | 100 |  |
|  | 3 | 100 |  |
|  | 4 | 100 |  |
|  | 5 | 100 |  |
|  | 6 | 100 |  |
|  | 7 | 100 |  |
|  | 8 | 100 |  |
|  | 9 | 100 |  |
| 2 | 1 | 50 | 60 |
|  | 2 | 65 | 70 |
|  | 3 | 45 | 60 |
| 4 | 1 | 40 | 60 |
|  | 2 | 60 | 70 |
|  | 3 | 50 | 70 |
| 6 | 1 | 50 | 50 |
|  | 2 | 70 | 60 |
|  | 3 | 40 | 70 |
| 8 | 1 | 45 | 70 |
|  | 2 | 70 | 60 |
|  | 3 | 35 | 60 |
| 10 | 1 | 55 | 60 |
|  | 2 | 50 | 70 |
|  | 3 | 45 | 60 |
| 12 | 1 | 55 | 60 |
|  | 2 | 55 | 50 |
|  | 3 | 30 | 50 |

