## Supplemental File Readme for "Transcriptomic and proteomic dynamics of ovarian follicle group culture resemble *in vivo* folliculogenesis"

**Supplemental File 1 (SFile_1):** Somatic cell and oocyte metadata containing ANOVA & clustering results

Tabs: SomaticCells (tab 1) and Oocytes (tab2)

Gene: ensemble id + '_' + gene symbol

[Day|Cond|Int]_pval: ANOVA p-values for given variables

[Day|Cond|Int]_fstat: ANOVA F-statistics for given variables

[Day|Cond|Int]_FDR: Benjamini-Hochberg corrected p-values from ANOVA

[Day|Cond|Int]_SOMcluster: SOM cluster assigned to variable-significant DEGs, N.S. = not significant.

Genes_for_phaseshift: genes kept for phase-shift analysis (Day- and Int-significant DEGs), following columns will be filled in only if TRUE

clusters_[5x|10x]_kmeans: kmeans cluster assigned to each condition for the given gene

same_kmeans: whether the kmeans clusters matched

Sin_[5x|10x]: phase shift curves assigned to condition

kmeans_delta: difference between the assigned kmeans clusters as numerics

kmeans_pair: clusters_5x_kmeans + '_' + clusters_10x_kmeans

kmeans_delta_label: label given to the kmeans_delta

Sin_delta: difference between the pahse shift curve assignments as numerics

Sin_pair: Sin_5x + '_' + Sin_10x

Sin_delta_label: label given to the Sin_delta

Sin_leading_split: sub-labels for Leading in Sin_delta_label

Sin_lagging_split: sub-labels for Lagging in Sin_delta_label

**Supplemental File 2 (SFile_2):** GO Biological Process for somatic cell Condition- and Interaction-significant DEG clusters.

Tabs: Condition (tab 1) and Interaction (tab 2)

Column 1/rownames: GO biological process name

num_of_genes: number of genes present in the pathway from reference list (all genes in our data)

[c0_0|c0_1|c1_0|c1_1]:FDR : FDR for the term in the given cluster

[c0_0|c0_1|c1_0|c1_1]:OR : odds ratio of the term in the given cluster.

**Supplemental File 3 (SFile_3):** GO Biological Process for somatic cells and oocytes phase curves for 5X and 10X

Tabs: SomaticCells (tab 1) and Oocytes (tab2)

Column 1/rownames: GO biological process name

num_of_genes: number of genes present in the pathway from reference list (all genes phase analysis)

top_hit_[5x|10x]: phase curve with lowest FDR for the term

top_fdr_[5x|10x]: lowest FDR for the term

[c1:10]:OR : odds ratio of the term in the given phase curve.

[c1:10]:FDR : FDR for the term in the given phase curve

**Supplemental File 4 (SFile_4):** GO Biological Process for somatic cells and oocytes phase curves for 5X and 10X for pathways that are significant in 5X and 10X

Tabs: SomaticCells (tab 1) and Oocytes (tab2)

Column 1/rownames: GO biological process name

num_of_genes: number of genes present in the pathway from reference list (all genes phase analysis)

top_hit_[5x|10x]: phase curve with lowest FDR for the term

top_fdr_[5x|10x]: lowest FDR for the term

delta: difference between the assigned phase curves

delta_lab: label for the delta

Highlighted in Fig [2|3]: “x” indicates is used in figures

**Supplemental File 5 (SFile_5)**: Ligand and receptor metadata

Tabs: four interactions

[cond|day|int]_fval: F-values from ANOVA for the three terms

[cond|day|int]_pval: p-values form the ANOVA for the three terms

[cond|day|int]_FDR: FDR from the ANOVA for the three terms

ct: cell type (Oocytes or cells)

type: type of interaction (self or directional)

gene: pair name

set.x: interaction pair (same as tab)
lig: Ligand name
rec: Receptor name

*following columns are only for L-R pairs used in phase analysis
[FiveX|TenX]_max_cor: the max correlation value with the 10 phase curves
[FiveX|TenX]_max_hit: the phase curve with the max correlation
[FiveX|TenX]_second_cor: the second max correlation value with the 10 phase curves
[FiveX|TenX]_second_hit: the phase curve with the second max correlation
Sine_pair: the phase curve assignment for the gene: 5X + '_' + 10X

**Supplemental File 6 (SFile_6)**: protein GO results
Tabs: Day-specific and Cluster GO terms

Name: GO Biological Process name
### Genes in GO Term: number of genes in the go term from the reference data (all proteins)
[id]:FDR : FDR for the cluster/day for given term
[id]:OR : odds ratio for the cluster/day for given term

**Supplemental File 7 (SFile_7)**: protein metadata
rownames: protein Accession ID
gene_name: Mouse gene name pulled from Accession ID
[Day|Cond]_fval: F-statistic from ANOVA
[Day|Cond]_FDR: FDR from ANOVA
cluster: cluster identity if Day_FDR < 0.10
